# Photopharmacology in Action: Conformational Landscape of a Photoswitchable Covalent Kinase Inhibitor

**DOI:** 10.1101/2025.05.27.656308

**Authors:** Thilo Duve, Sebastian Thallmair

## Abstract

Photopharmacology is a rapidly evolving field that uses light to control drug activity with high spatial and temporal precision, offering innovative therapeutic strategies with reduced side effects. In this study, we investigate a photoswitchable covalent inhibitor for the MAP kinase JNK3, a target for the treatment of neurodegenerative diseases. The inhibitor, which is based on a diazocine photoswitch, can undergo reversible photoisomerization of a double bond, switching between two (meta-)stable isomers. Atomistic molecular dynamics simulations, comprising almost 40 µs of total simulation time, reveal how the distinct conformational spaces of the two isomers modulate their interactions with JNK3 within the ATP-binding pocket. We show that only the metastable *trans* isomer can form a covalent bond with JNK3, thereby permanently inhibiting its function. In contrast, for the *cis* state the distance between the inhibitor and the targeted cysteine residue is too large to allow covalent bond formation. Furthermore, our simulations reveal that the covalent bond, in combination with the environment of the protein pocket, hinders the full back-relaxation of the *trans* isomer to the stable *cis* isomer. Finally, our data shows that the covalently bound stable *cis* isomer modulates the conformations of JNK3, mainly of its activation loop. Our findings provide molecular insights into the complex dynamics of photoswitchable inhibitors, guiding future drug design for light-controlled therapeutics.

## Introduction

Molecular photoswitches are capable of undergoing reversible three-dimensional structural changes upon irradiation with light, typically via isomerization of a double bond or bond formation or cleavage [1, 2]. The prototypical photoswitch applied in many studies is azobenzene, which switches via an isomerization of its central N=N double bond [3, 4]. This study focuses on diazocine, a bridged azobenzene derivative that serves as a photoswitch. Unlike the closely related azobenzene, where the *trans* isomer is thermodynamically more stable, diazocine is known to preferentially adopt the *cis* isomer. Consequently, photoisomerization leads to a structural transition from a bent (*cis* isomer) to a more stretched geometry (*trans* isomer) [5, 6]. This is particularly relevant for biological applications, where the extended isomer often resembles the active molecule. Thus, the use of diazocine instead of azobenzene can result in thermodynamically stable molecules which are inactive in the dark that can be activated via irradiation — a strategy known as *sign inversion* [6]. Diazocine displays well-separated absorption bands for its isomers, facilitating efficient and reversible photo-switching. Photoisomerization from the stable *cis* isomer to the metastable *trans* isomer is induced by blue light (*λ ≈* 370 − 400 nm). Back-conversion (*trans* to *cis*) can be triggered either by green light (*λ ≈* 480 − 550 nm) or via thermal relaxation over a timescale ranging from minutes to hours [5, 7, 8]. The light-induced reversible structural changes of photoswitches make them highly attractive for diverse applications in material science [9–12], energy storage [13], biological systems [12, 14, 15], and beyond. In particular, their integration into pharmaceutical molecules offers innovative approaches for spatial and temporal control of the bioactivity of compounds. Photopharmacology, an emerging interdisciplinary field, exploits such photoswitchable compounds to modulate drug function with high spatio-temporal precision [4, 16].

This work investigates a diazocine-based, photoswitchable JNK3 inhibitor previously reported by Reynders et al. [17]. The MAP kinase c-Jun N-terminal kinase 3 (JNK3) is a key player in neuronal cell biology and contributes to neuronal differentiation [18]. Compared to its ubiquitously expressed isoforms JNK1 and JNK2, the expression of JNK3 is limited to the brain, heart, and testes [18, 19], making it an attractive drug target [20]. JNK3 is involved in key mechanisms of neurodegenerative diseases, including Alzheimer’s, Huntington’s, and Parkinson’s disease [19, 20].

The photoswitchable covalent inhibitor (Figure 1A) targets the cysteine residue Cys154 located near the ATP-binding site of JNK3. The inhibitor consists of a pyridinylimidazole-based ATP-mimetic moiety that enables non-covalent binding to the ATP-binding pocket, connected via a diazocine photoswitch to a cysteine-reactive acrylamide warhead [17]. The proposed mechanism of the inhibitor is shown in Figure 1B. If it binds to the ATP-binding pocket in the less active *cis* isomer (red), the reactive group is positioned away from the targeted cysteine residue. Upon photoisomerization to the *trans* isomer (blue), the acrylamide warhead moves closely to the reactive cysteine, enabling covalent bond formation and subsequent inhibition of kinase activity. It was shown experimentally that in the dark, where the *cis* isomer dominates, it is a weak inhibitor. Blue-light irradiation (*λ* = 400 nm) induces a switch to the *trans* isomer, leading to a substantial increase in kinase activity inhibition [17].Back-relaxation of the photoswitchable inhibitor was achieved with green light (*λ ≈* 500 − 600 nm) [17].

**Figure 1.**
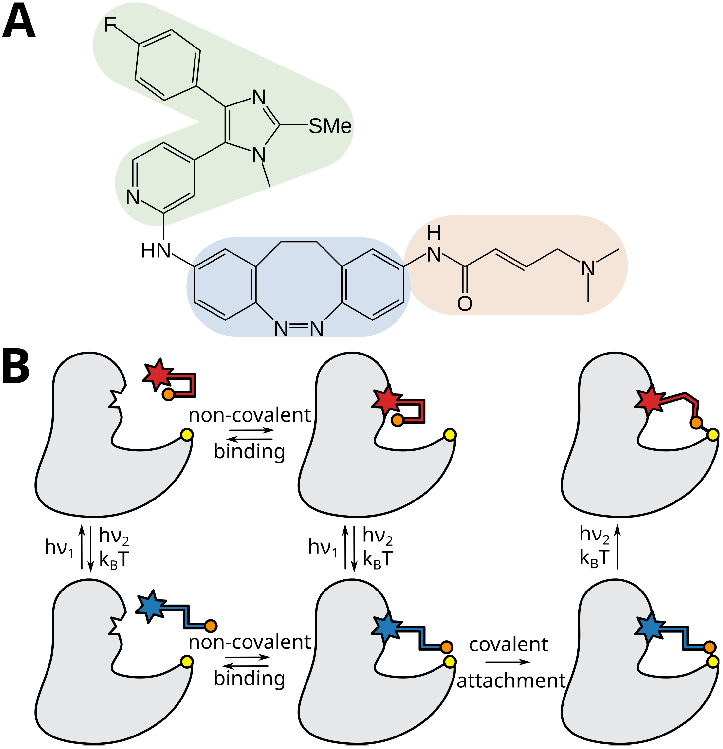
(A) Chemical structure of the photoswitchable JNK3 inhibitor [17]. The ATP-mimicking motif (green), diazocine photoswitch (blue) and reactive tail (orange) are highlighted. (B) Schematic illustration of the possible mechanisms of a photoswitchable covalent kinase inhibitor. The less active *cis* isomer is indicated in red, the more active *trans* isomer in blue. The reactive group of the photo-switchable inhibitor is shown in orange, the targeted cysteine residue is shown in yellow.

The conformational landscape of the diazocine photoswitch, compromised of six distinct conformations shown in Figures 2 and 4, introduces an additional layer of complexity to the dynamics and intermolecular interactions of the photoswitchable inhibitor. Therefore, one needs to determine which conformations are capable of binding non-covalently to the ATP-binding pocket of JNK3 and, subsequently, forming a covalent bond with the targeted cysteine residue, Cys154. In addition, the structural constraints imposed by the covalent attachment must be taken into account – in particular, whether the ATP-binding pocket permits the back-isomerization of the photoswitch or if this process can only occur upon dissociation of the inhibitor from the ATP-binding pocket of JNK3.

**Figure 2.**
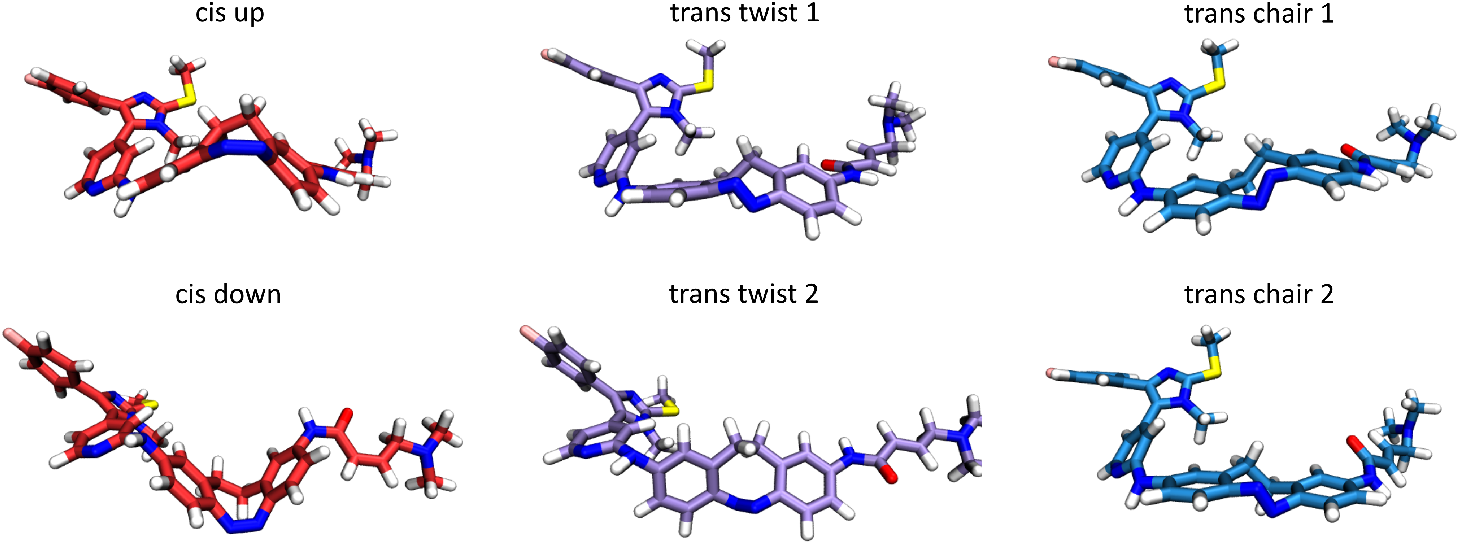
Snapshots of the photoswitchable inhibitor in the six distinct conformations after equilibration.

In this study, we investigate the dynamical interactions between the photoswitchable inhibitor and JNK3 using atomistic molecular dynamics (MD) simulations. We first characterize the conformational landscape of the diazocine photoswitch and the diazocine-based JNK3 inhibitor. Then, we examine the complex dynamics of the photoswitchable inhibitor within the ATP-binding pocket of JNK3, highlighting how both the binding site environment and the covalent attachment to the protein modulate the inhibitor’s conformational flexibility in its *cis* and *trans* isomeric states. Our analysis reveals that the targeted cysteine residue is significantly more accessible to the *trans* isomer, supporting the proposed mechanism of covalent inhibition. Finally, we investigate how inhibitor binding influences the conformational dynamics of JNK3 itself.

## Computational Methods

### Computational Details

Semiempirical quantum mechanical MD simulations used to validate the parametrization of diazocine were performed using the GFN2-xtb implementation in the software package Bartender [21, 22]. 100 ns long simulations of diazocine where performed with an implicit solvent model for water in the in the *cis* up, *trans* chair 1 and *trans* twist 1 conformations, respectively [21, 23]. Simulations of the *cis* up, *trans* chair 2, and *trans* twist 2 conformations were not performed as they are chemically identical to their respective pseudo-enantiomer.

All classical MD simulations in this work were performed using GROMACS 2020.5 [24, 25] with the CHARMM36m force field [26]. The pressure of the system was controlled using the Berendsen barostat [27] for equilibration simulations and the Parrinello-Rahman barostat [28] for all production runs. All simulations used an isotropic pressure coupling with a reference pressure of 1.0 bar; the compressibility was set to 4.5 *×* 10^−5^ bar^−1^ (*τ*_*p*_ = 5.0 *ps*). A Nose-Hoover thermostat [29] was used to maintain the temperature. Constraints were solved using LINCS, with a LINCS order of 4 in a single iteration (lincs_iter = 1). Electrostatic interactions were calculated with the smooth particle mesh Ewald (PME) method [30]. The minimization and equilibration of the system followed the standard CHARMM-GUI solvation builder procedure [31, 32]. All systems were minimized using the steepest-descent algorithm, followed by an equilibration run for 125 ps in the *NpT* ensemble at 303.15 K with a time step of 2 fs. The production run was conducted again in *NpT* ensemble at 303.15 K (*τ*_*t*_ = 1.0 *ps*) with a time step of 2 fs. For simulations of diazocine in water, one repeat of 100 ns was performed. For simulations of the photoswitchable inhibitor in water, one repeat with a simulation time of 1 µs was performed. For all other systems, three repeats with a respective simulation time of 1 µs were performed. A detailed overview of the simulations performed in this work is shown in Table 1. Overall, 39.6 µs of simulations were performed and analyzed.

**Table 1:** Listing of the total simulation time of this work.

| Conformation | Diazocine | Water | Pocket | Pocket + Bond | Sum |
| --- | --- | --- | --- | --- | --- |
| *Cis* down | $2 \times 0.1 \mu\text{s}^1$ | $1 \times 1.0 \mu\text{s}$ | $3 \times 1.0 \mu\text{s}$ | $3 \times 1.0 \mu\text{s}$ | 7.2 $\mu\text{s}$ |
| *Cis* up | $1 \times 0.1 \mu\text{s}$ | $1 \times 1.0 \mu\text{s}$ | $3 \times 1.0 \mu\text{s}$ | $3 \times 1.0 \mu\text{s}$ | 7.1 $\mu\text{s}$ |
| *Trans* chair 1 | $2 \times 0.1 \mu\text{s}^1$ | $1 \times 1.0 \mu\text{s}$ | $3 \times 1.0 \mu\text{s}$ | $3 \times 1.0 \mu\text{s}$ | 7.2 $\mu\text{s}$ |
| *Trans* chair 2 | $1 \times 0.1 \mu\text{s}$ | $1 \times 1.0 \mu\text{s}$ | $3 \times 1.0 \mu\text{s}$ | — | 4.1 $\mu\text{s}$ |
| *Trans* twist 1 | $2 \times 0.1 \mu\text{s}^1$ | $1 \times 1.0 \mu\text{s}$ | $3 \times 1.0 \mu\text{s}$ | $3 \times 1.0 \mu\text{s}$ | 7.2 $\mu\text{s}$ |
| *Trans* twist 2 | $1 \times 0.1 \mu\text{s}$ | $1 \times 1.0 \mu\text{s}$ | $3 \times 1.0 \mu\text{s}$ | — | 4.1 $\mu\text{s}$ |
| JNK3 apo | — | $3 \times 1.0 \mu\text{s}$ | — | — | 3.0 $\mu\text{s}$ |
| **Sum** | 0.9 $\mu\text{s}$ | 9.0 $\mu\text{s}$ | 18.0 $\mu\text{s}$ | 12.0 $\mu\text{s}$ | 39.9 $\mu\text{s}$ |

<sup>1</sup> Including one replica of semiempirical QM calculations using GFN2-xtb [21].

### System Setup and Small Molecule Parametrization

Initial topologies and parameters for the diazocine photoswitch and the photoswitchable inhibitor were obtained using CHARMM-GUI’s Ligand Reader & Modeler based on the CHARMM General Force Field (CGenFF) [31, 33], resulting in a single topology for each molecule that supports all six conformations, depending on the starting structure. The missing residues 373-379 in the protein structure of JNK3 were modeled with the SWISS-MODEL web tool [34] using the crystal structure of the protein (PDB-code: 7ORE [17]) and the amino acid sequence of JNK3 as a reference. Terminal missing residues were not modeled, resulting in a structure covering the structured domain of JNK3 with the residues 44-402 as shown in Figure 3.

**Figure 3.**
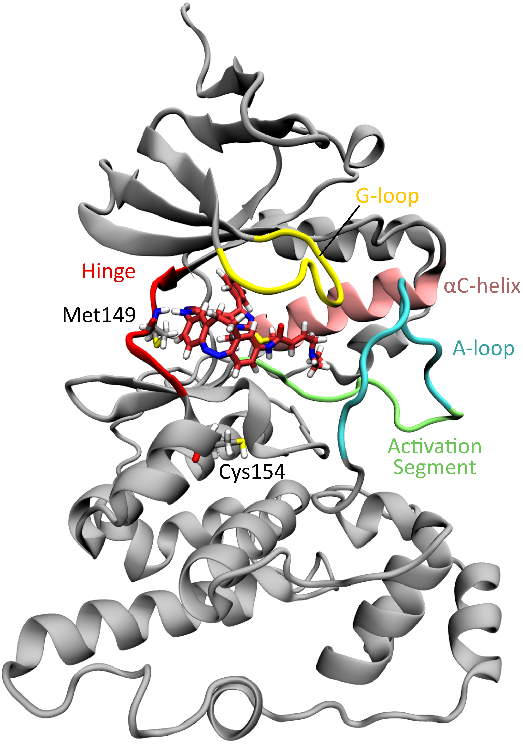
Three-dimensional structure of JNK3. The photoswitchable inhibitor in the *cis* down conformation bound non-covalently to the ATP-binding pocket is shown in red licorice. The residues Met149, whose backbone forms two hydrogen bonds with the photoswitchable inhibitor, and Cys154 targeted for covalent bond formation are shown in licorice. The following regions are highlighted by color: G-loop (residues 71-78, yellow), *α*C-helix (residues 102-117, salmon), hinge region (residues 147-152, red), A-loop (residues 217-226, cyan), and the activation segment, containing the A-loop (residues 207-226, lime).

Simulation systems and topologies were generated with the CHARMM-GUI Solution Builder [31, 32]. Structures were placed in a cubic box of 10.0*×*10.0*×*10.0 nm^3^ for simulations containing the protein and 4.5 *×* 4.5 *×* 4.5 nm^3^ for simulations of small molecules. The systems were solvated using the TIP3P water model [35]. For simulations containing JNK3, NaCl was added at a concentration of 150 mM, and the system was neutralized by adding counter ions. Starting structures for the *trans* chair 1 & 2 and *trans* twist 1 & 2 conformations of diazocine and the photoswitchable inhibitor were obtained by manually modifying the positions of the atoms in the azo-group of the photoswitch. It was ensured that the diazocine unit adopts the respective conformation during the minimization and equilibration of the system as seen in Figure 2.

To accurately represent the covalently bound form of the photoswitchable inhibitor, it was reparameterized to account for the removal of the double bond in the acrylamide group. In the process, two hydrogen atoms were added to the C_*α*_ and C_*β*_-atoms. To avoid a renumbering of the system, a dummy particle type without van der Waals interactions and charge was created. This type was then assigned to the hydrogen atom attached to the sulfur of the cysteine residue that would relo-cate during the bond formation, as well as the hydrogen atom attached to the neighboring C_*α*_-atom of the inhibitor that was added previously. The replacement of a regular atom with a dummy particle prevents interactions with other atoms, essentially removing it from the simulation, without a change of the number of atoms in the system’s topology [36, 37]. To retain an integer charge for the system, the negligible partial charge of both hydrogen atoms was dissipated to a methyl group distant from the binding pocket. This was done to minimize the influence of the partial charge on the system. The covalent bond between the photoswitchable inhibitor and JNK3 was added using the [`intermolecular_interations`] block in the topology file of the system. The reference bond distance of 1.8 Å was measured in the crystal structure of the inhibitor bound to JNK3 [17]. The force constant (1.66 *×* 10^5^ kJ mol^−1^nm^−2^) is the standard force constant for a sulfur-carbon bond in the CHARMM36m force field.

An alternative approach to model the covalently bound inhibitor is the recently released CHARMM-GUI PDB Reader for covalent ligand modelling [38]. A comparison between the simulations of this work and the potentials generated by the CHARMM-GUI tool is shown in Figure S6. The inclusion of these parameters could yield improved ligand conformations.

### Analysis

The protein backbone root mean square fluctuation (RMSF) was calculated using an in-house script that can be found at https://github.com/cristina-gil-herrero/RMSFscript. Trajectories were divided into 200 ns segments, and the protein was centered in the simulation box. In seven iterations, the trajectory was fitted to all C_*α*_ atoms that exhibited a mean RMSF value below the threshold of 1.5 Å in the previous iteration by using *gmx trjconv* with the flag *-fit rot+trans*. This was done to avoid fitting the trajectory to highly flexible regions of the protein. The final mean RMSF values for each residue were calculated following the seventh iteration. RMSF values were computed using *gmx rmsf* with the flags *-nofit -res*. The mean RMSF of the C_*α*_ atom *i* was calculated according to:

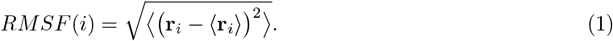

Additional analysis was conducted using the *MDAnalysis* Python package v2.8.0 [39, 40]. Atomic distances, angles, and dihedral angles were calculated using *MDAnalysis* fast distance array computation functions. Principle component analysis (PCA) was performed using *MDAnalysis*.*analysis*.*pca*. The respective trajectories were then projected on the first component using *pca*.*transform* and the following equation:

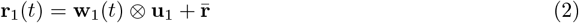

Where **r**_1_(*t*) is the vector of coordinates in the projected trajectory, **w**_1_(*t*) are the weights of the first principle component, **u**_1_ the eigenvector and 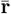 the mean atom positions. The root mean square deviation (RMSD) of the inhibitor was calculated using *MDAnalysis*.*analysis*.*rms* according to Equation 3, which calculates the Euclidean distance between the current configuration and the reference configuration, with *N* being the number of coordinates [41, 42]. Trajectories were fitted to the C_*α*_ atoms within 5 Å of the photoswitchable inhibitor in the crystal structure (residues 33, 34, 36, 38, 39, 41, 43,54-56, 87, 89, 107-118, 121, 155-159, 169, 189, 191, 192, and 224).

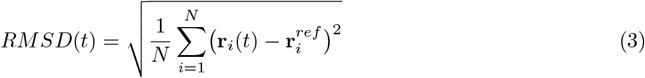

Figures were prepared with the Python libraries Matplotlib v.3.10.0 [43] and Seaborn v0.13.2 [44]. Renders of molecules were prepared using the program packages Visual Molecular Dynamics (VMD) version 1.9.4 [45] and PyMOL version 3.1 [46]. Chemical structures were created using ChemDraw 2016.

## Results and Discussion

Diazocine exists in two isomers – the thermodynamically stable *cis* isomer and the metastable *trans* isomer. The chemical structure of diazocine is shown in Figure 4A. The *cis* isomer shows two boat-like conformations, which interconvert via boat inversion. This interconversion is slow on the NMR time scale at room temperature [8, 47]. The *trans* isomer shows four distinct conformations, two twist and two chair conformations, with the twist being the more stable and predominant conformation at room temperature [7, 8, 48]. All four conformations can interconvert into each other and are at equilibrium at room temperature. In total, six conformations exist for diazocine (Figure 4B) [5]. Because of the low equilibration rate of the isomers, no interconversion is to be expected on the time scale of atomistic MD simulations. We thus consider them as distinct conformations. Moreover, when asymmetric substitution patterns are present on the aromatic rings of diazocine, the similar conformations are mirror images of each other. Therefore, we will use the term pseudo-enantiomers in the following for conformation pairs, such as *cis* up and *cis* down, or *trans* chair 1 and *trans* chair 2.

**Figure 4.**
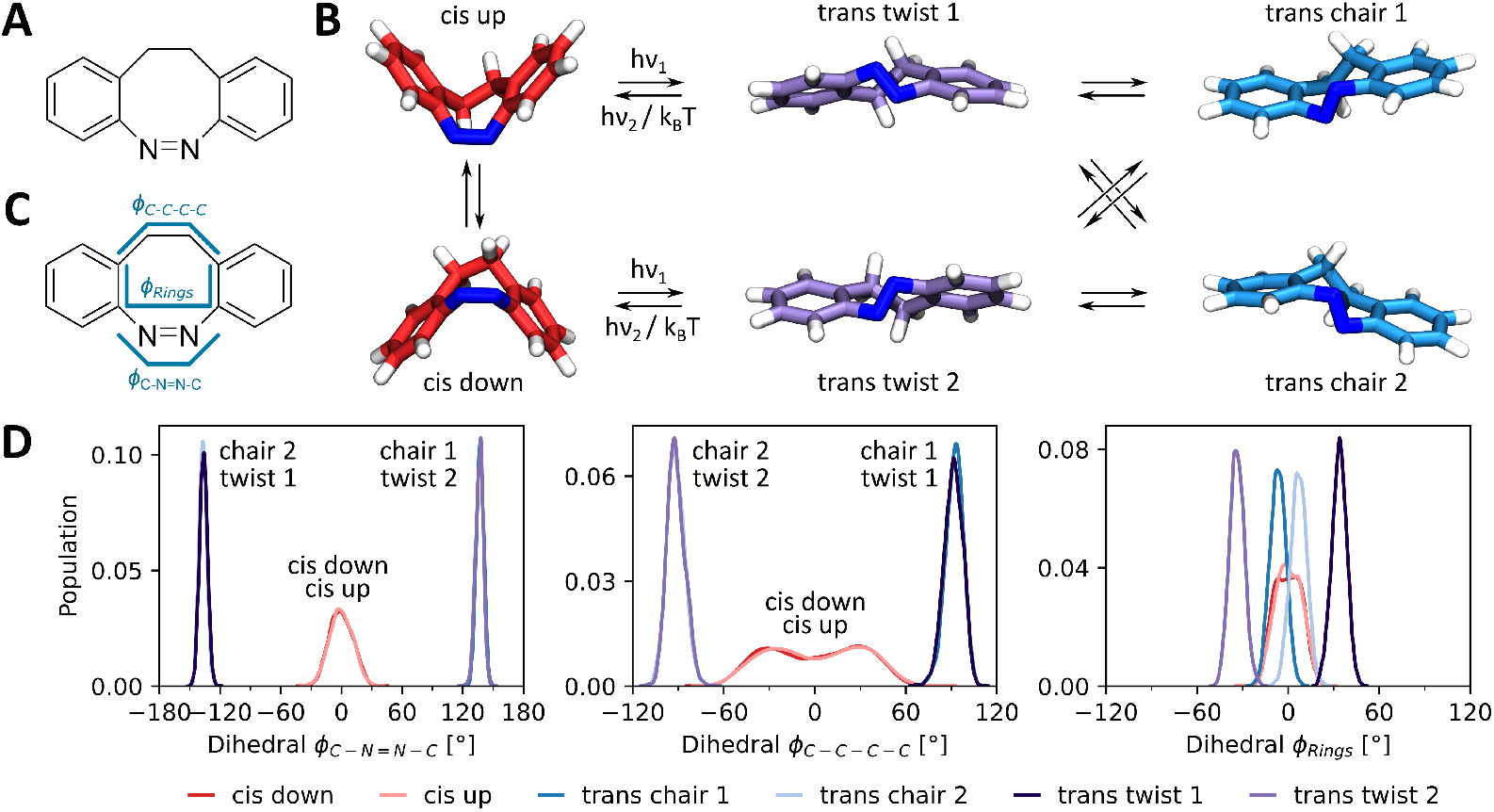
(A) Chemical structure of the diazocine photoswitch. (B) Configurational and conformational space of diazocine. (C) Dihedral angles analyzed to identify the diazocine conformations. (D) Distributions of the characteristic dihedral angles for all conformations of diazocine.

The CHARMM36m topology of diazocine supports all six conformations of diazocine, however, no interconversion is observed, and the obtained conformation is solely determined by the starting structure. In this study, we do not focus on the photo-induced isomerization process of diazocine, which occurs on a timescale of 30 to 500 fs [49]. Instead, we aim to characterize the dynamics of the photoswitchable inhibitor in the electronic ground state in its six distinct conformations before and after photoswitching on the nanosecond to microsecond timescale. Thus, all simulations in this work focus on one of the six conformations of the diazocine photoswitch separately.

To characterize the conformational space of diazocine, MD simulations of 100 ns were performed for each of the six conformations in water. While we note the limitations associated with the use of a single replica and the short simulation time, the small size of the molecule suggests that the conformational space should be appropriately sampled within this timescale. The configuration of diazocine can be described by three characteristic dihedral angles that are depicted in Figure 4C with the distributions throughout the 100 ns trajectories depicted in Figure 4D. Additionally, these dihedral angles were validated by comparing the distributions to semi-empirical QM simulations. Figure S2 shows a reasonable overlap between the QM and atomistic distributions.

The dihedral angle *ϕ*_C-N=N-C_ describes the state of the azo group and is found to be 0 ± 11^°^ for the *cis* isomer and ±137 ± 4^°^ for the *trans* isomer. The dihedral angle *ϕ*_C-C-C-C_, which represents the state of the aliphatic C_2_H_4_ bridge, exhibits a broad bimodal distribution centered at 0^°^(3 ± 26^°^) for the *cis* isomer, while the *trans* isomer adopts a value of ±93 ± 5^°^. Looking at the four conformations of the *trans* isomer, one can see that for the *trans* chair conformations, the azo group and the aliphatic bridge are oriented parallel to each other, resulting in distributions for *ϕ*_C-N=N-C_ and *ϕ*_C-C-C-C_ with identical signs (positive for chair 1 and negative for chair 2). In the *trans* twist conformations, the azo group and the aliphatic bridge have a perpendicular orientation, resulting in inverted signs for the dihedral angle distributions (Figure 4B, D). The relative orientation of the benzene ring edges in the eight-membered central ring, described by the dihedral angle *ϕ*_Rings_, varies between different c onformations. I n the *cis* and *trans* chair conformations, the benzene ring edges remain approximately parallel, whereas the *trans* twist conformations exhibit a characteristic dihedral angle of ±35 ± 5^°^, which gives this conformation its name.

With the complex conformational space of the diazocine photoswitch characterized, the next objective was to determine how the various conformations influence the i nteraction o f the diazocine-based photoswitchable inhibitor (Figure 1A) with its target protein, the protein kinase JNK3 [17]. To investigate this, we conducted extensive MD simulations of the inhibitor in aqueous solution and within the ATP-binding pocket of JNK3, accumulating a total simulation time of almost 40 µs (Table 1). For this, the photoswitchable inhibitor with the diazocine photoswitch in one of the six conformations described previously was aligned with the crystal structure [17]. The simulation system is depicted in Figure 5. Across all simulations, no unbinding events were observed throughout the simulations, and the inhibitor’s binding pose remained stable for the 1 µs simulation time of the individual replicas. The ATP-binding motif closely matched the binding pose observed in the crystal structure (Figure S3). As experimentally determined in the crystal structure, this binding pose is stabilized via two hydrogen bonds with the backbone of Met149 in the hinge region of JNK3 (Figure S4). Additionally, the binding pose is stabilized via a hydrogen bond between the protonated amino group of LYS93 and the nitrogen in the imidazol ring of the photoswitchable inhibitor, an interaction not present in the crystal structure (Figure S4). Thus, our simulations suggest a strong interaction between the inhibitor and JNK3 on the short microsecond timescale.

**Figure 5.**
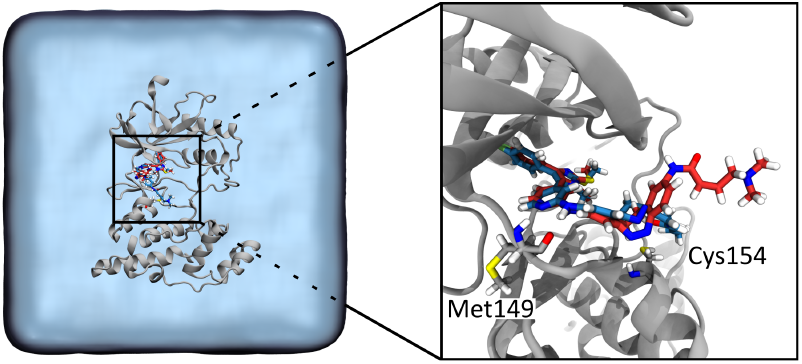
Setup of the simulation box. Starting structures are based on the crystal structure of JNK3 (silver, PDB-code: 7ORE [17]) and the photoswitchable inhibitor. The targeted cysteine residue Cys154 and residue Met149 are indicated. The system is solvated in water (blue transparent surface) and 150 mM NaCl (not shown).

The inhibition efficiency of the photoswitchable inhibitor is determined by its ability to form a covalent bond with the targeted cysteine residue Cys154 following the non-covalent binding to JNK3 [50]. The reaction mechanism for this covalent bond formation is shown in Figure 6A. To estimate the inhibition potency of the isomers, the probability of covalent bond formation between the photo-switchable inhibitor and JNK3 was estimated by examining the accessibility of Cys154 to the reactive acrylamide group of the inhibitor. No other cysteine residue is located within reach of the reactive group of the photoswitchable inhibitor (Figure S5). Since conventional MD simulations do not capture chemical reactions, covalent bond formation could not be directly studied. Instead, two important metrics for the formation of the transition state (Figure 6A) were employed to estimate the likelihood of covalent bond formation: (i) the distance between the atoms forming the covalent bond (*r*_*S,Cβ*_) and(ii)the angle of attack between the cysteine residue and the reactive acrylamide (*α*_*S,Cβ*_,*C*_*α*_) indicated in Figure 6A. The results of this analysis are shown in Figure 6B, while the time series of *r*_*S,Cβ*_ and *α*_*S,Cβ*_,*C*_*α*_ are shown in Figures S7, and S8.

**Figure 6.**
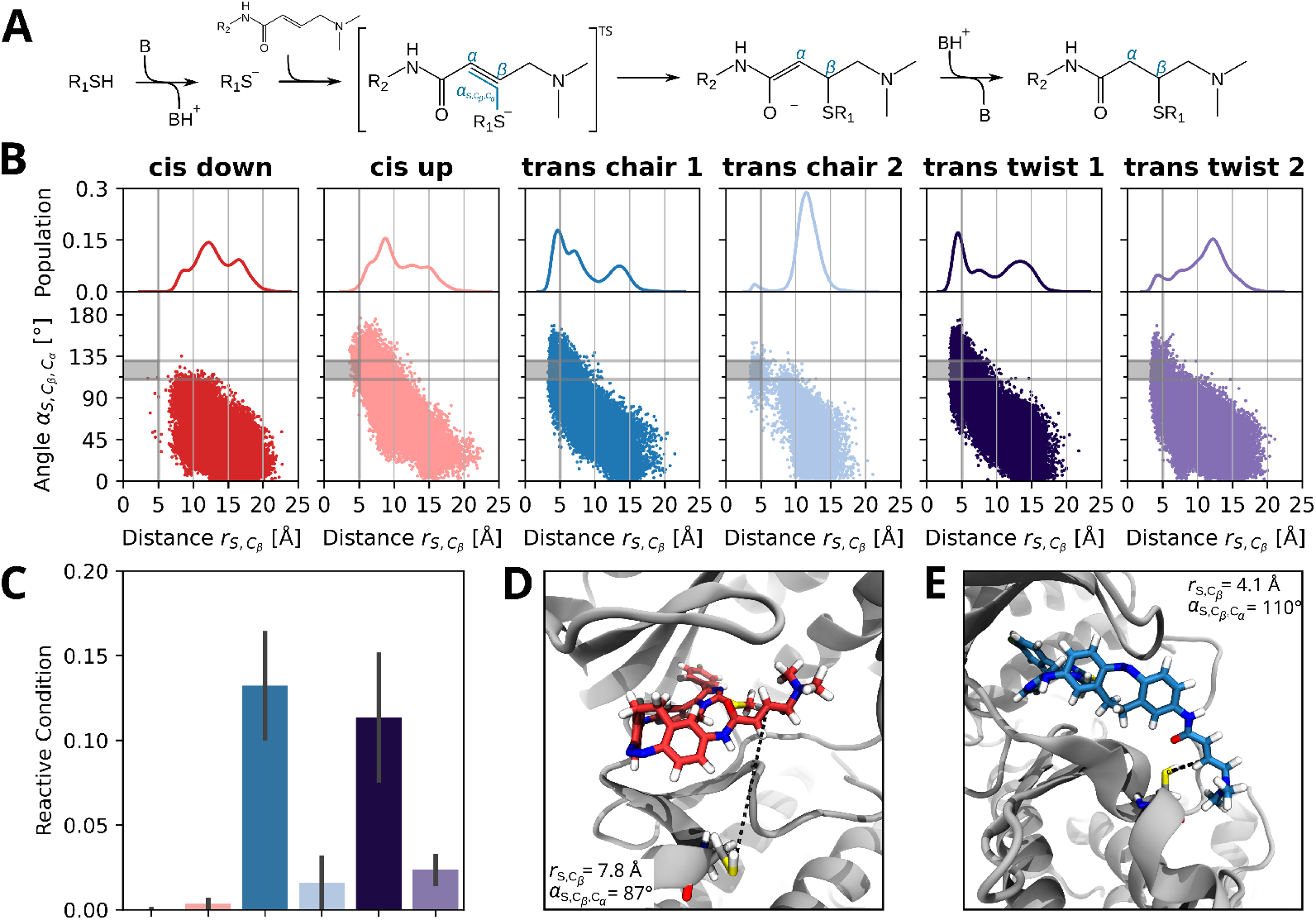
(A) Reaction mechanism of thiol addition to the acrylamide group of the photoswitchable inhibitor. C_*α*_ and C_*β*_ atoms and the angle of attack *α*_*S,Cβ*_,*C*_*α*_ are indicated in blue. TS indicates a transition state. (B) Distribution of distances between sulfur atom of Cys154 and C_*β*_ of the photo-switchable inhibitor (*r*_*S,Cβ*_) and scatter plot of the distance and the angle of attack *α*_*S,Cβ*_,*C*_*α*_. The reactive condition is indicated by a grey rectangle. (C) Average number of contacts allowing bond formation between Cys154 and the photoswitchable inhibitor, defined as the fraction of frames with *r*_*S,Cβ*_ *≤* 5.0 Å and 110^°^ *≤ α*_*S,Cβ*_,*C*_*α*_ *≤* 130^°^ corresponding to the rectangle indicated in (B). Error bars represent the standard error of the mean over 200 ns windows. (D), (E) Representative snapshots of the photoswitchable inhibitor non-covalently bound to the ATP-binding pocket in the *cis* down (D) and *trans* chair 1 (E) isomer.

The distance analysis of *r*_*S,Cβ*_ is limited by the lack of deprotonation of the sulfur atom in the cysteine residue, an essential step in the mechanism of covalent bond formation [51]. Furthermore, the distance between the two atoms is inherently limited by the Van der Waals radii of the atoms involved, which do not account for the possibility of the formation of a transition state. Both of these factors lead to a lower bound of the distance. In our simulations, a minimum distance of 3.3 Å was reached. Due to this, the distance should only be viewed as an estimate of the probability of bond formation. The time-evolution of the distance *r*_*S,Cβ*_ is shown in Figure S7. The angle of attack (*α*_*S,Cβ*_,*C*_*α*_, Figure 6A), an important property of the transition state, was analyzed to refine the estimation of the likelihood of bond formation. Literature values for transition states indicate angles between 110^°^ and 112^°^ [52]. In our simulations of the covalently bound inhibitor, observed angles ranged from 110^°^ to 130^°^. Based on these findings, as an estimation, cutoffs of *r*_*S,Cβ*_ *≤* 5.0 Å and 110^°^ *≤ α*_*S,Cβ*_,*C*_*α*_ *≤* 130^°^ were used to determine the reactive condition. Figure 6C depicts the relative number of frames at which the reactive condition is met. It clearly shows that the *trans* chair 1 and *trans* twist 1 conformations have the highest probability for covalent bond formation. For the *cis* isomer the probability is negligible. Figure S9in the SI shows that the observed trends are independent of the exact choice of the cutoffs. Both conformations of the *cis* isomer – *cis* down and *cis* up – adopt a similar geometry, with the kink of the photoswitch pointing toward the ATP-pocket of JNK3. This orientation results from a rotation around the inhibitor’s nitrogen hinge. However, differences arise due to the positioning of the reactive tail. For the *cis* down conformation, the peptide group of the acrylamide tail is oriented away from Cys154, while it is oriented towards Cys154 for its pseudo-enantiomers, the *cis* up conformation (Figure S11). This leads to the *cis* up conformation showing a higher population below 7.5 Å. However, neither conformation shows relevant populations below the cutoff of 5.0 Å. The angle of attack of the *cis* down conformation is also low, with most values being below 100^°^, while the *cis* up conformation spans from 0 to 180^°^, with a sizable population in the targeted range between 110^°^ and 130^°^. In our simulation the *cis* isomer of the photoswitchable inhibitor fulfills the reactive condition in <1% of the frames, and therefore, the probability of covalent bond formation is very low for both *cis* conformations.

This behavior is consistent throughout all three repeats as shown in Figure S10. For the *trans* isomer, the different conformations exhibit notable differences. Both the *trans* chair 2 and *trans* twist 2 conformations show a small population below 5.0 Å, with a considerably larger population centered at 12 Å. The latter conformation is stabilized by a hydrogen bond between the carboxyl-oxygen atom of the peptide group and the backbone of Ser72 in the G-loop, where the reactive group is oriented away from Cys154, leading to the increased distance (Figure S11, S12). Similar to the *cis* isomer, both the *trans* chair 2 and *trans* twist 2 conformation show a low probability of covalent bond formation, with 1.6% and 2.4% of frames fulfilling the reactive condition, respectively.

In contrast, the respective pseudo-enantiomers *trans* chair 1 and *trans* twist 1 routinely achieve lower distances, coupled with attack angles within the transition state range. Both conformations exhibit three populations at distances of approximately 4.8, 6.0, and 13.5 Å. The lowest-distance population displays angles of attack between 60 and 160^°^. Comparing *trans* chair 1 and *trans* twist 1, the relative weight of these populations differs: while both conformations predominantly occupy the lowest distance population, the 6.0 Å population is more prevalent for *trans* chair 1 than for twist 1. The opposite is the case for the population centered at 13.5 Å. The *trans* chair 1 and *trans* twist 1 conformations show a significantly higher probability of covalent bond formation, with 13% and 11% of frames fulfilling the reactive condition. While the *trans* twist 1 conformation shows high variability between repeats, with the reactive condition being met predominantly in repeat 1, the behavior of *trans* chair 1 conformation is more consistent in all three repeats (Figure S10). The photoswitchable inhibitor is brought into position for the reactive condition via a hydrogen bond between the side chain of ASN152 and the peptide carbonyl group of the photoswitchable inhibitor. Figure S15 shows that for the *trans* isomer the reactive condition is met if the hydrogen bond is formed, indicating a key mediating role of ASN152 in covalent bond formation. Additionally, ASN152 could facilitate the covalent bond forming reaction by stabilizing the negatively charged intermediate via the hydrogen bond (Figure 6A).

Despite the limitations of MD simulations in analyzing the probability of covalent bond formation, a difference between the distance and angle of attack distributions of the *cis* and *trans* isomers was observed. The results suggest that the probability of covalent bond formation is higher for *trans* chair 1 and *trans* twist 1 compared to the *cis* isomer and the *trans* chair 2 and twist 2 conformations. While we did not observe any differences for the non-covalent binding of the *cis* and *trans* isomers, our results suggest that the overall equilibrium of the *trans* isomer is shifted towards the bound inhibited state, as covalent bond formation depopulates the non-covalently bound state, which is repopulated from the free state (Figure 1B). These findings align with experimental data that the photoswitchable inhibitor acts as a weak inhibitor of JNK3 in the dark (IC_50_ = 507 nM, dominated by the *cis* isomer) but exhibits significantly stronger inhibition after pulse irradiation (IC_50_ = 31.9 nM, dominated by the *trans* isomer) [17].

We assessed the conformation of the elongated, covalently bound crystal structure of the photoswitchable inhibitor (PDB-code: 7ORE [17]) by analyzing the characteristic dihedral angles of diazocine discussed previously (Figure 4). The aliphatic C_2_H_4_-bridge exhibits a dihedral angle of *ϕ*_*C*−*C*−*C*−*C*_ = 107.7^°^, which, as expected, falls within the distribution of the *trans* isomer, specifically the chair 1 and twist 2 conformations. However, the azo group, with a dihedral angle of *ϕ*_*C*−*N*=*N*−*C*_ = 1.8^°^, does not align with the *trans* isomer distribution but instead falls in the *cis* isomer’s distributions (*cis* down and *cis* up).

Simulations of diazocine and the photoswitchable inhibitor starting from the elongated crystal structure in aqueous solution revealed that diazocine adopts the *cis* down conformation during equilibration and remains in this state throughout the production runs of 0.1 µs and 1.0 µs, respectively. These results confirm that – as expected – the azo group primarily dictates the state of the photoswitch, rather than the aliphatic bridge. This finding further validates the parametrization of diazocine and the photoswitchable inhibitor, demonstrating its capability to reproduce the expected conformational behavior. Based on these observations, we hypothesize that the elongated crystal structure represents a partially relaxed state, in which the azo group has relaxed back to the thermodynamically stable, less active *cis* isomer, while the aliphatic C_2_H_4_-bridge and the reactive tail remain in the active state. The latter two have higher flexibility and thus adapt to the constraints due to the covalent bond between Cys154 and the inhibitor. The reported crystal structure might have resulted from an overlap of the *trans* with the *cis* conformation, as partial relaxation of the sample might have occurred during data collection. Therefore, it remains open if the photoswitch state in the *trans* labeled crystal structure represents the only conformation or if there are further conformations hidden behind the broadly spread electron density shown in Figure S13.

The covalently bound crystal structure serves as an ideal starting point to study the effects of back-relaxation. As shown in the previous section, the inhibitor forms a covalent bond in the metastable *trans* isomer. However, the photoswitch relaxes thermally to the thermodynamically stable *cis* isomer within minutes to hours [5, 7, 17]. To examine the behavior of the covalently bound photoswitchable inhibitor before and after back-relaxation, simulations were initiated for the inhibitor in the *cis* down, *cis* up, *trans* chair 1, and *trans* twist 1 conformations. No simulations were conducted for chair 2 and twist 2, as these conformations are not expected to form a covalent bond with high probability (Figure 6C, S9), nor is relaxation to these states expected, unlike for the *cis* conformations. The simulations for the *cis* down, *trans* chair 1, and twist 1 conformations were initiated from the crystal structure (manually modifying the azo-group position where necessary), while the *cis* up conformation was started from a snapshot of simulations of the non-covalently bound inhibitor, where the reactive condition was fulfilled. Then the covalent bond was introduced before the start of the simulation. The different starting structure for the simulation of the *cis* up conformation results in a different protein conformation as shown Figure S14.

Our simulations show that the binding pose of *trans* isomer is stabilized by the previously discussed hydrogen bond between the side chain of ASN152 and the peptide carbonyl (Figure S16). While the *cis* up conformation’s binding position is stabilized by a less frequent hydrogen bond with the side chain of SER193 (Figure S17), the *cis* down conformations forms no hydrogen bonds outside of the previously discussed hydrogen bonds with the ATP-mimicking motif.

Again, the characteristic dihedral angles of the photoswitch were analyzed. In addition, the end-to-end photoswitch distance was calculated. This distance was defined between the two aromatic C atoms, which are bound to the ATP-mimicking motif, and the acrylamide tail, respectively (Figure S1). The *trans* isomer in the chair 1 and twist 1 conformations exhibited identical conformational distributions across all conditions – in water, non-covalently bound, and covalently bound to JNK3 – indicating that the addition of the covalent bond does not restrict its conformational space (Figures S18 and S19). This further stresses the ability of the *trans* isomer to form the covalent bond, as the resulting conformation is native to the *trans* state of diazocine without imposing significant conformational strain on the molecule.

In contrast, the combination of the covalent bond and the non-covalent binding in the ATP-binding pocket significantly restricted the conformational space of the *cis* isomer (Figure 7). Structural changes were observed when analyzing the photoswitchable inhibitor after back-relaxation, starting from the elongated crystal structure and transitioning into the *cis* down conformation. The dihedral angle *ϕ*_*C*−*N*=*N*−*C*_ exhibited a shift towards positive values, increasing from −0±11^°^ to 10±9^°^. Furthermore, the flexibility of the aliphatic C_2_H_4_-bridge was significantly reduced, transitioning from a bimodal distribution in the absence of the covalent bond (−6 ± 33^°^) to a unimodal distribution centered at −44 ± 16^°^, with a considerably narrower spread. Additionally, a twist in the diazocine photoswitch emerged, resembling the *trans* twist 2 conformation, as indicated by the dihedral angle *ϕ*_Rings_. The end-to-end distance also increased from 6.1 ± 0.3 Å in water to 6.6 ± 0.3 Å.

**Figure 7.**
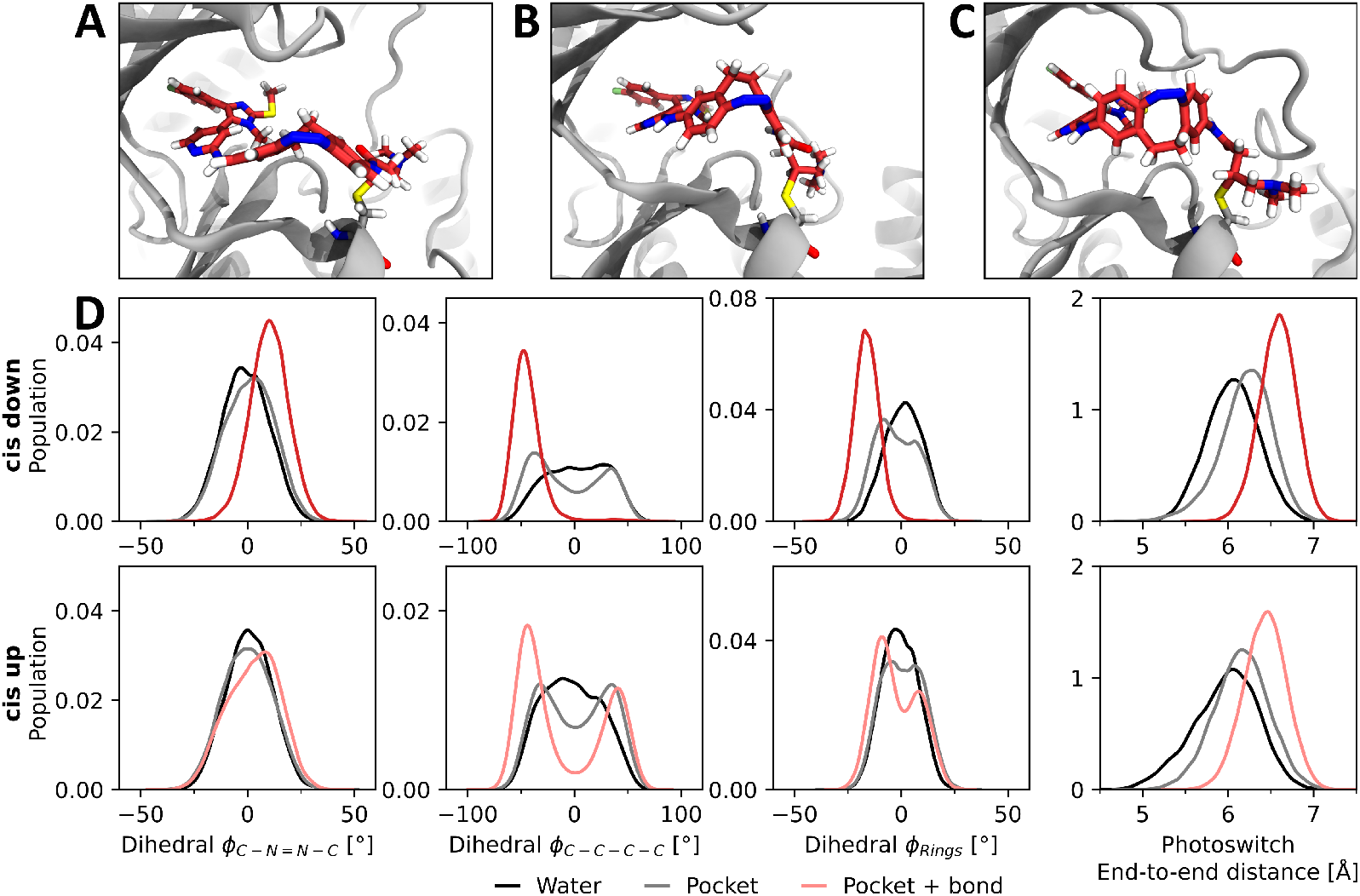
(A) Elongated crystal structure of the photoswitchable inhibitor used to study the covalently bound inhibitor after back-relaxation. (B), (C) Representative snapshots of the photoswitchable inhibitor covalently bound to the ATP-binding pocket in the *cis* down (B) and *cis* up (C) isomer. (D) Distributions of characteristic photoswitch dihedral angles and end-to-end distance for three systems– the photoswitchable inhibitor (*cis* up and *cis* down) in water, bound non-covalently (pocket) and covalently (pocket + bond) to JNK3. The dihedral angles and the end-to-end distance analyzed are indicated in Figure 4C and Figure S1, respectively.

These results suggest that full back-relaxation is not possible due to conformational constraints imposed by the covalent bond and the non-covalent binding to the ATP-binding pocket. Furthermore, these restrictions affect the hydrogen bond with the backbone NH of Met149, leading to increased distances between the interacting atoms (Figure S4).

Even though simulations for the covalently bound inhibitor in the *cis* up conformation were started from a native conformation of the inhibitor, the conformational space of the inhibitor is restrained, and the photoswitch is stretched, similar to the *cis* down isomer. While *ϕ*_*C−N*=*N−C*_ remains unchanged, *ϕ*_*C−C−C−C*_ and *ϕ*_Rings_ both favor negative angles over positive angles. In addition, the end-to-end distance increases from 6.0 ± 0.4 Å in water to 6.4 ± 0.3 Å and the ATP-mimicking motif of the photoswitchable inhibitor slightly changes its position, while the hydrogen bonds stay intact (Figure S4). This leads to increased RMSD values compared to the non-covalently bound inhibitor, as can be seen in Figure S3.

The simulations provide key insights into the conformational behavior of the covalently bound photoswitchable inhibitor before and after back-relaxation. While the *trans* isomer remains largely unaffected by the formation of the covalent bond, preserving its native conformational space, the *cis* isomer experiences significant structural constraints due to the covalent bond and the ATP-binding pocket. Throughout the simulations, the covalently bound inhibitor never left the ATP-binding pocket, although it could be speculated that the conformational stress in the back-relaxed *cis* isomer might pull the inhibitor out of the ATP-binding pocket of JNK3. The maintained binding mode in the ATP binding pocket is in agreement with experimental results, which show that the inhibition of JNK3 is permanent and cannot be reverted by back-relaxation to the *cis* isomer, once the covalent attachment took place [17].

So far, we have focused on the behavior of the photoswitchable inhibitor and how its conformations are influenced by the protein environment. However, the inhibitor might also impact the conformation of JNK3. It has been shown that the activation loop adopts distinct conformations which are linked to the activity of JNK3 [53]. To assess the influence of the photoswitchable inhibitor and its different conformations on the conformation of JNK3, we analyzed the dynamics of the hinge region and activation loop (see Figure 3). Both the hinge and A-loop can respectively adopt two distinct states, an open and a closed state. The conformation of these regions of JNK3 is closely linked to the inhibition/activation of the kinase activity. The closed state of the hinge and the A-loop open state have been linked to JNK3 inhibition [53, 54].

All simulations started from JNK3 co-crystallized with the photoswitchable inhibitor or a simulation snapshot with the photoswitchable inhibitor in the ATP-binding pocket and therefore in the hinge-open state. Throughout the simulations, no transition from the hinge-open to the hinge-closed state was observed in either the inhibitor-bound state or apo JNK3. In the inhibitor-bound state, this transition was not expected, as the protein was anticipated to behave similarly to ATP-bound JNK3, where hinge closure has not been observed [53]. Moreover, hinge closure occurs on a 2 µs timescale, whereas our simulation time was 1 µs per replica. Additionally, Lys43, a key residue in the intermediate state of the hinge closure, was absent from our simulations due to it not being resolved in the crystal structure used here [17, 53]. This, along with the limited simulation time, likely explains the absence of hinge closure in apo JNK3.

Further analysis focused on the A-loop within the activation segment (residues 207–226), which can adopt an open and closed conformation, respectively (Figure S20). To quantify the A-loop conformation we performed a principle component analysis (PCA). The first principle component correlates with the movement of the A-loop as shown in Figure S22. The histogram of the first principle component (Figure S23) and visual inspection as shown in Figure 8 show that for apo JNK3, the A-loop samples both open and closed states with a preference for the open state. This is in agreement with previous computational works [53].

**Figure 8.**
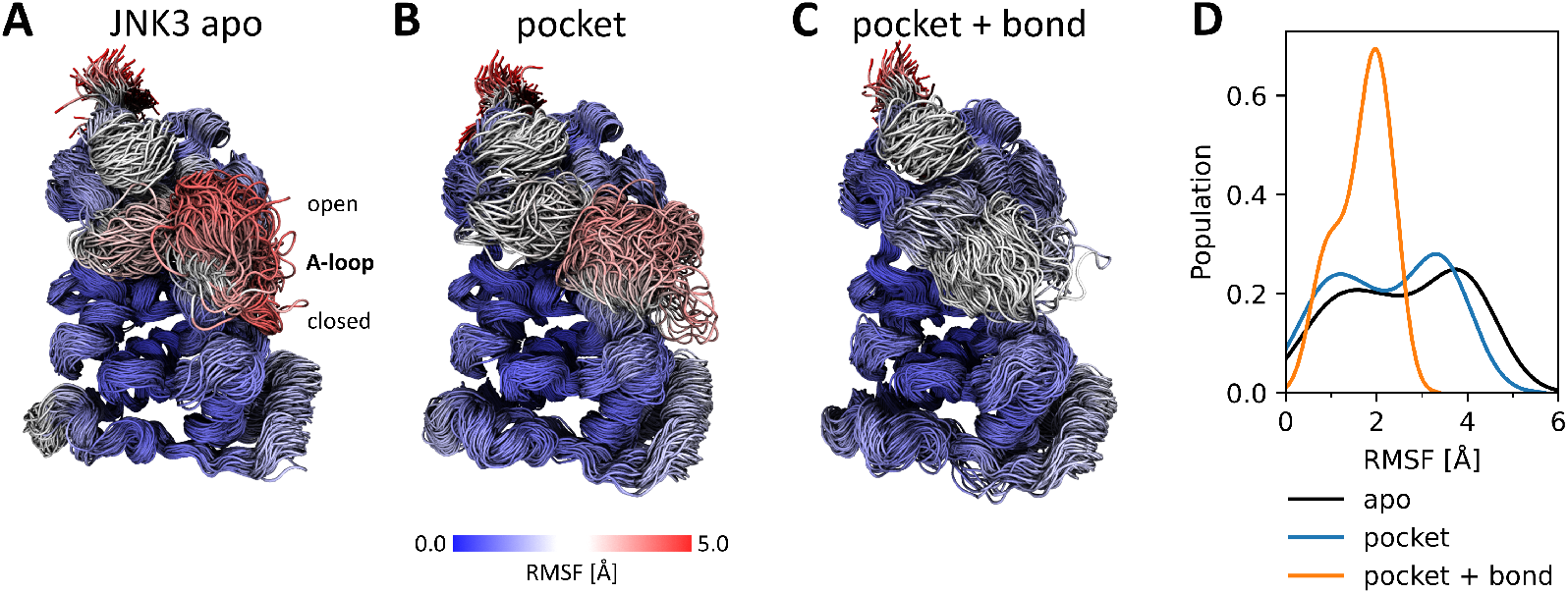
Protein conformations of JNK3 in the apo state (A) and with the photoswitchable inhibitor in the *cis* up isomer bound non-covalently (B) and covalently (C) to the ATP-binding pocket. Indicated is the conformation of the A-loop in the open and closed states. 300 frames across 3 repeats of 1 µs each are shown, the *C*_*α*_ RMSF with the respective inhibitor binding is indicated by the color. (D) Distribution of the backbone (C_*α*_) RMSF of the activation segment of JNK3 (residues 207-226) in the apo state and with the photoswitchable inhibitor in the *cis* up conformation bound non-covalently and covalently to the ATP-binding pocket of JNK3.

For simulations including the photoswitchable inhibitor, the A-loop remained predominantly in the closed state both for the non-covalently and covalently bound inhibitor. This observation is in agreement with previous computational works that showed this behavior for simulations of ATP bound to JNK3. This shows that the photoswitchable ligand has a similar impact on the protein conformation of JNK3 as ATP. One exception to this is the photoswitchable inhibitor in the *cis* up conformation. In simulations with the non-covalently bound inhibitor the A-loop samples both states with a slight preference for the open state. In simulations with the covalently bound inhibitor in the *cis* up conformation only the closed state is observed, as can be seen in the first principle component of the PCA (Figures S22, S23. This also leads to a reduced flexibility of the A-loop as quantified by the root-mean-square fluctuation (RMSF) of the C_*α*_-atoms. Note that in the case of the covalently bound *cis* up conformation, the analysis might be biased by the different protein conformation in the starting structure as shown in Figure S14. For the other simulations no significant changes in protein flexibility were observed (Figures S24, S25).

These findings suggest that the switch state of the photoswitchable inhibitor influences the protein conformation. The strong preference of the A-loop for the open state in the covalently bound, back-relaxed *cis* up isomer, may explain the experimentally observed irreversibility of JNK3 inhibition [17].

In this conformation, the open A-loop could obstruct the ATP-binding pocket, preventing inhibitor dissociation and additionally stabilizing the inhibited state. However, it should be noted that the simulation timescale of 1 µs limits the extent to which long-term conformational changes in JNK3 can be assessed.

## Conclusions

In summary, our MD simulations, spanning nearly 40 µs, provide a thorough characterization of the conformational dynamics of a photoswitchable JNK3 inhibitor. The results demonstrate that the distinct conformations of the diazocine photoswitch isomers significantly influence the interactions between the photoswitchable inhibitor and the protein. These findings offer a structural explanation for the differences in inhibitory strength observed between the two isomers [17]. Furthermore, our simulations reveal that the constraints imposed on the inhibitor by the ATP-binding pocket of JNK3, along with the covalent bond, hinder full back-relaxation to the stable *cis* isomer and induce conformational stress on the inhibitor. Despite this conformational stress, we could identify several aspects supporting the experimentally observed irreversibility of JNK3 inhibition by the photoswitchable inhibitor following irradiation and potential covalent attachment. Notably, the covalently attached inhibitor was not observed to dissociate from the pocket, even after back-relaxation, suggesting that the conformational stress exerted by the covalent bond and the ATP-binding pocket is not sufficient to remove the inhibitor from the ATP-binding pocket on the low microsecond timescale. Additionally, the conformational changes of JNK3 observed for the covalently bound inhibitor of the *cis* isomer may contribute to the stabilization of the inhibited state, with the preference of the A-loop for the open state potentially preventing inhibitor dissociation.

Taken together, our results demonstrate the utility of MD simulations in providing insights into the molecular mechanisms by which photoswitchable compounds interact with their target proteins and modulate their activity. The observed conformational dynamics underscore the importance of accounting for the photoswitch behavior beyond a simplistic view of the switch state during the design of such inhibitors.

A limitation of this study is the automatized parametrization of the photoswitchable inhibitor. Future studies could use QM methods to further optimize the bonded parameters and provide insight in the electronic structure of the different conformation of the inhibitor and whether all conformations can be represented with the same point charges.

Nevertheless, whether the photoswitch’s state directly affects the inhibitor’s non-covalent binding affinity to JNK3 remains an open question. Future studies employing coarse-grained molecular dynamics simulations may offer further insights into the non-covalent binding mechanisms of these systems [55, 56]. Furthermore, our simulations could serve as a starting point to investigate the covalent bond formation between the photoswitchable inhibitor and JNK3 by QM/MM simulations [57] as well as to study the photo-induced *cis-trans* isomerization of the inhibitor by QM/MM or nuclear quantum dynamics simulations [58].

## Supporting Information

Zenodo repository https://doi.org/10.5281/zenodo.17600146. Molecular topologies, simulation parameters, starting structures and structures of representative binding poses.

Supporting Information. Supplementary Figures on photoswitchable inhibitor conformation, characterization of inhibitor binding pose, estimation of covalent bond formation probability, behavior of covalently bound inhibitor, and impact of photoswitchable inhibitor on the conformation of JNK3.

## Acknowledgements

The authors thank Cristina Gil Herrero for critically reading the manuscript, Saara Lautala for technical assistance, and Stefan Knapp for fruitful discussions. Furthermore, we thank the Alfons und Gertrud Kassel Foundation, the Dr. Rolf M. Schwiete Foundation, the H. & E. Kleber Foundation, the SCALE cluster of excellence initiative, and the Center for Multiscale Modelling in Life Sciences (CMMS) sponsored by the Hessian Ministry of Science and Art for funding and the Center for Scientific Computing at Goethe University Frankfurt for access to Goethe-HLR.

## Table of Content Graphic

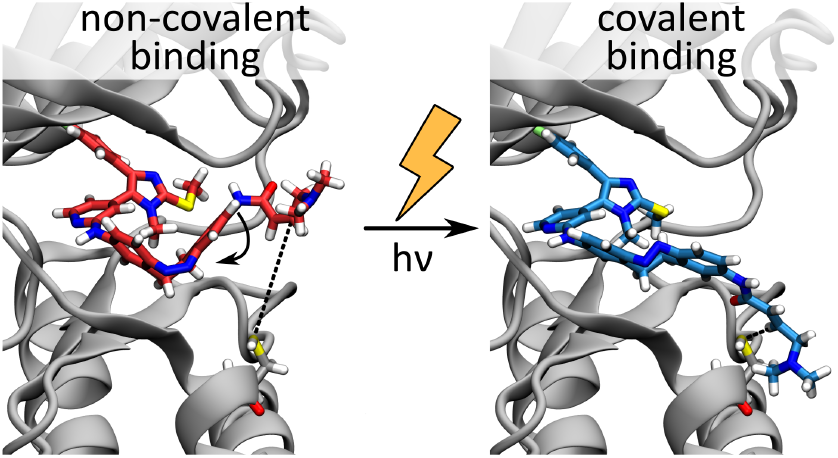

