## Supporting Information for "Photopharmacology in Action: Conformational Landscape of a Photoswitchable Covalent Kinase Inhibitor"

### Contents

S1 Photoswitch End-to-End distance . . . . . . . . . . S2
S2 Validation of diazocine parametrization . . . . . . . . . . S2
S3 Stability of the inhibitor binding pose . . . . . . . . . . S3
S4 Estimation of the covalent bond formation probability . . . . . . . . . . S4
S5 Behavior of the photoswitchable inhibitor bound covalently to JNK3 . . . . . . . . . . S9
S6 Impact of the photoswitchable inhibitor on JNK3 conformation . . . . . . . . . . S14

### S1 Photoswitch End-to-End distance

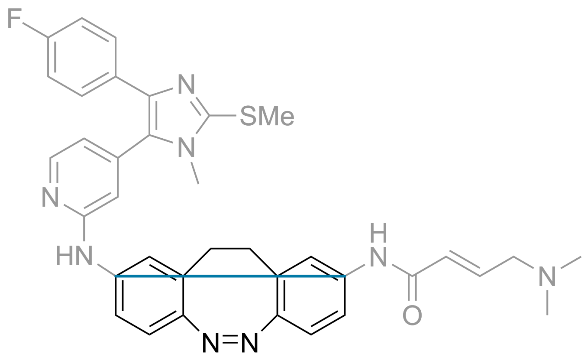


Figure S1: End-to-end distance of the photoswitch (indicated in blue).

### S2 Validation of diazocine parametrization

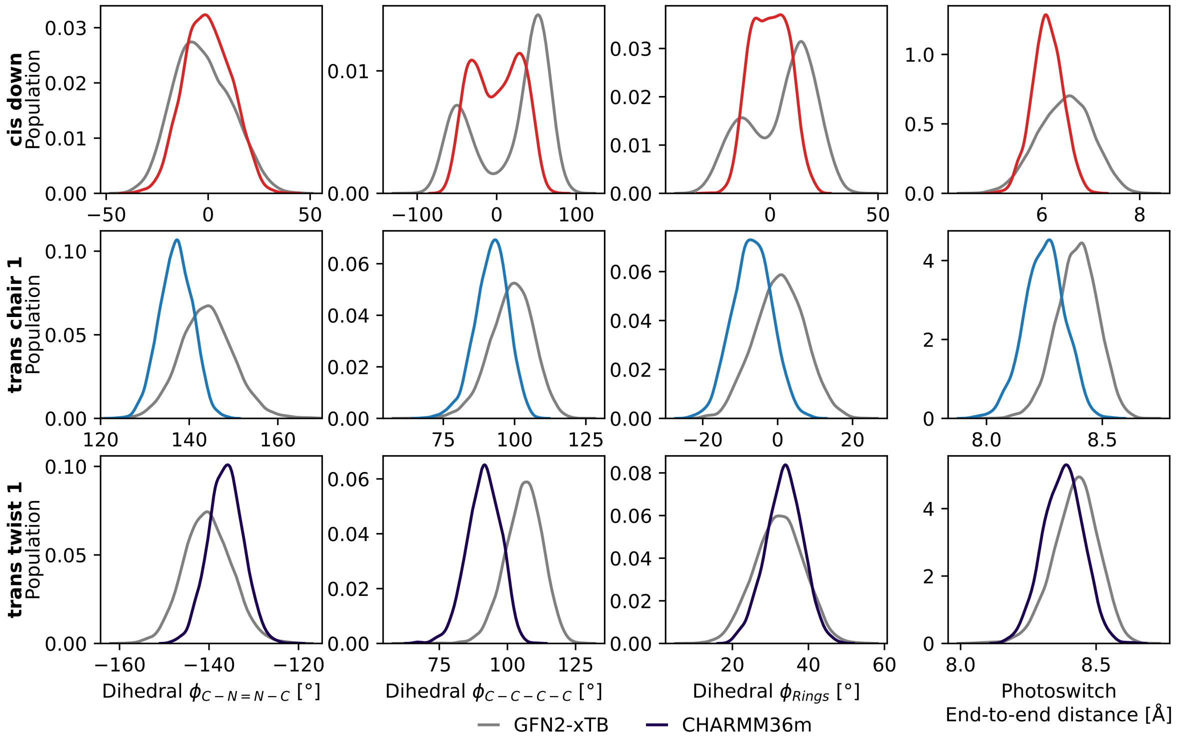


Figure S2: Distributions of the characteristic photoswitch dihedral angles and end-to-end distance of diazocine, simulated with GFN2-xtb (semiempirical quantum mechanical level) and CHARMM36m (atomistic MD). The dihedral angles are indicated in Figure 3 and the distance in Figure S1.

### S3 Stability of the inhibitor binding pose

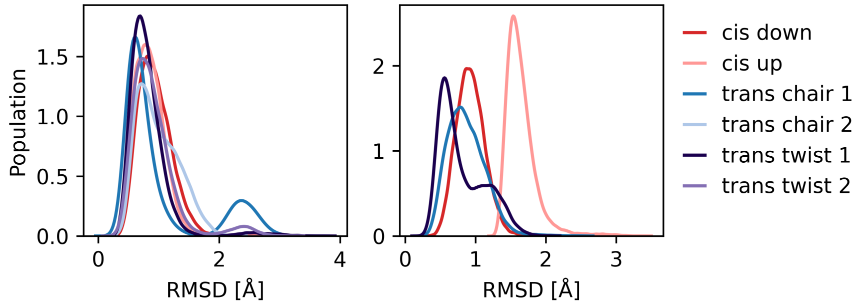


Figure S3: Inhibitor RMSD of ATP-mimicking motif relative to the crystal structure of the photoswitchable inhibitor in the ATP-binding pocket of JNK3. Trajectories were fitted to $C_\alpha$ atoms within 5 Å of the photoswitchable inhibitor in the crystal structure.

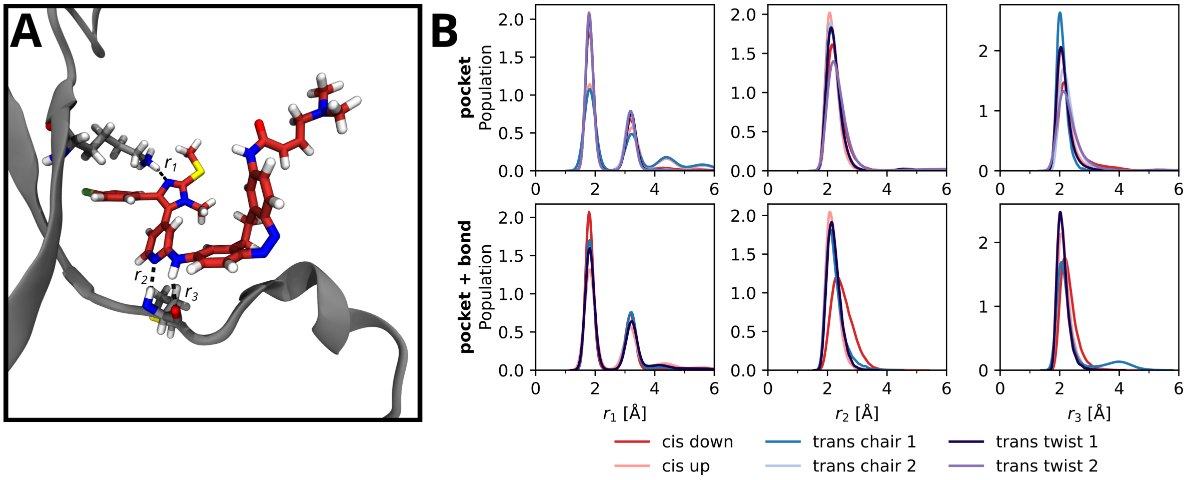


Figure S4: Hydrogen bonds between the ATP-mimicking motif of the photoswitchable inhibitor and JNK3. (A) Snapshot of the photoswitchable inhibitor in the ATP-binding pocket of JNK3. Indicated are the distances for the hydrogen bonds with LYS93 ($r_1$), the backbone aminogroup ($r_2$), and the backbone carbonyl of MET149 ($r_3$). (B) Distributions of $r_1$, $r_2$, and $r_3$ in the simulations of the photoswitchable inhibitor bound non-covalently (pocket) and covalently (pocket + bond) to JNK.

### S4 Estimation of the covalent bond formation probability

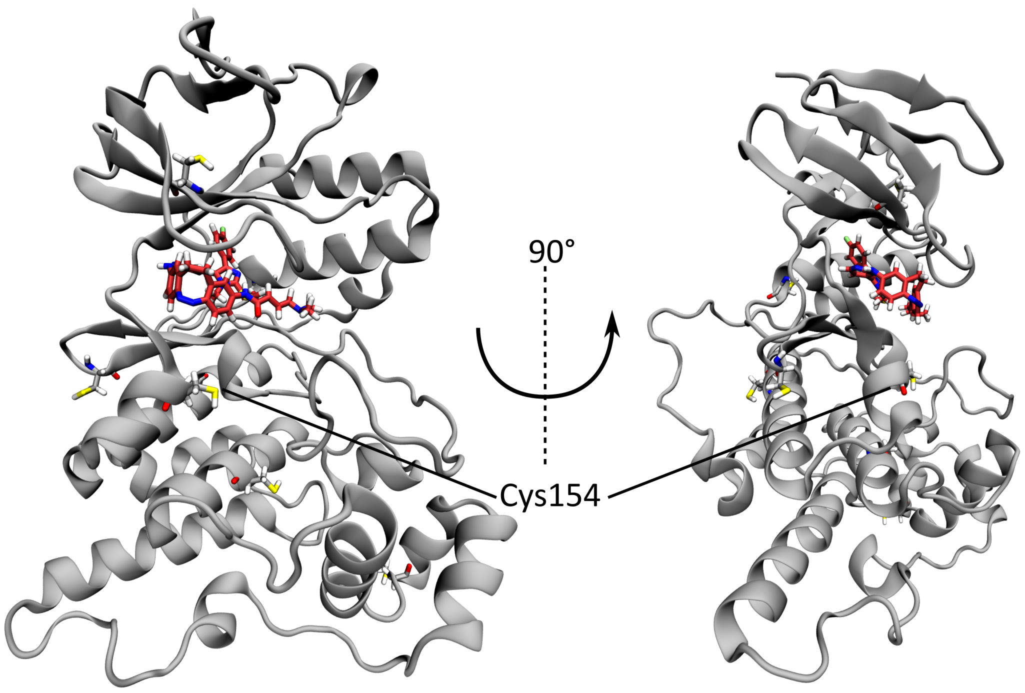


Figure S5: All cysteine residues of JNK3, shown in licorice (residues 79, 117, 154, 175, 201, 251, and 283). Residue Cys154, targeted for covalent bond formation, is highlighted. It is the only cysteine residue in reach of the photoswitchable inhibitor when bound non-covalently to the ATP-binding pocket of JNK3.

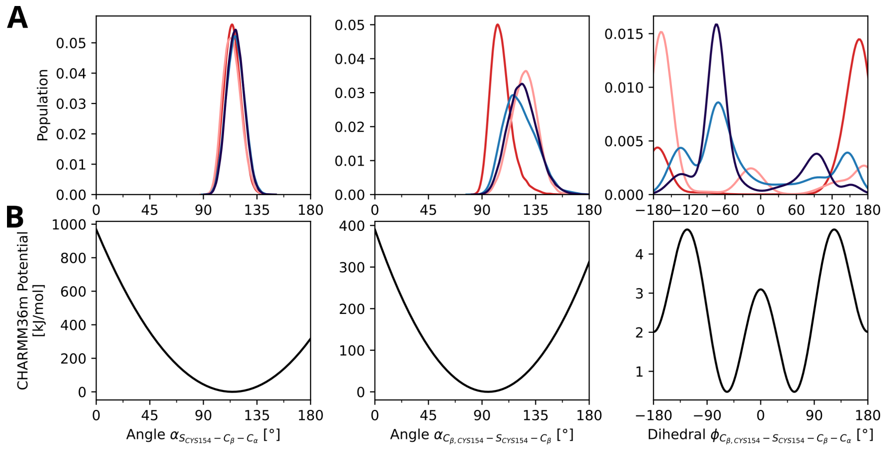


Figure S6: Comparison of angles and dihedrals around the covalent bond between the simulations in this work and the CHARMM-GUI PDB Reader for covalent ligand modelling [1]. (A) Distributions of two angles and a dihedral along the covalent bond between CYS154 and the photoswitchable inhibitor. (B) Bonded potentials generates by the CHARMM-GUI PDB Reader for covalent ligand modelling for the angles and dihedral mentioned previously.

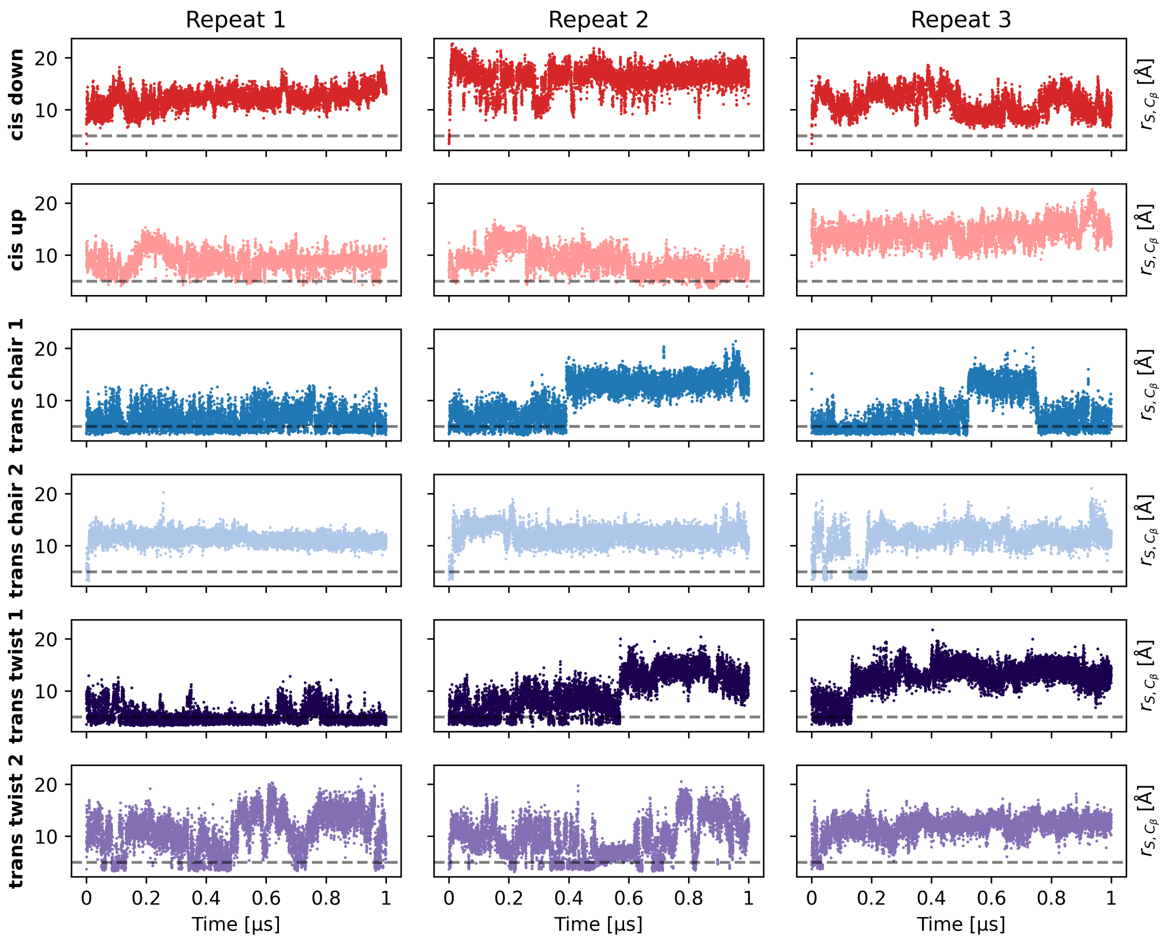


Figure S7: Time series for the distance between the (non-covalently bound) photoswitchable inhibitor and the sulfur atom of Cys154 ($r_{S,C_\beta}$). The dashed line indicates a cutoff of 5.0 Å used to distinguish between reactive ($r_{S,C_\beta} \leq 5.0$ Å) and non-reactive ($r_{S,C_\beta} > 5.0$ Å) conformations.

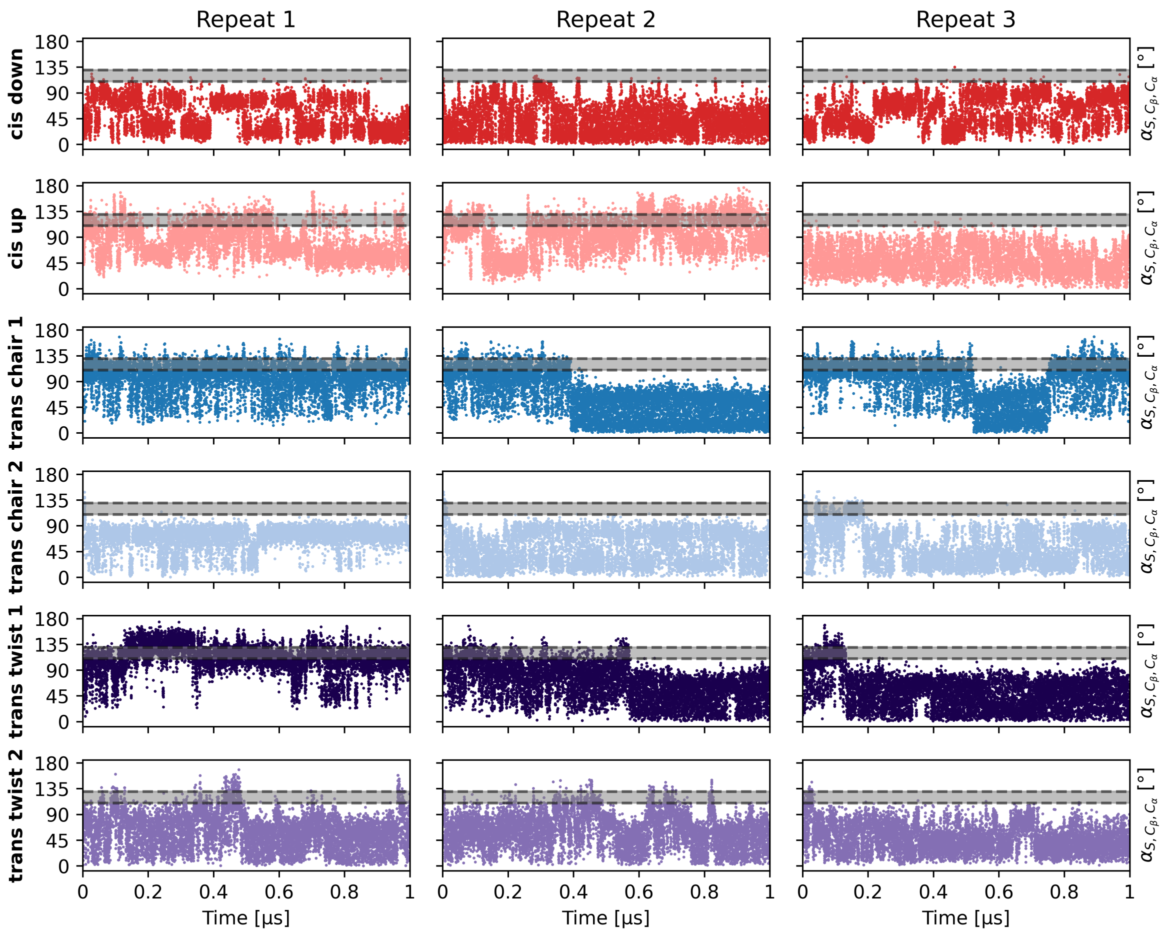


Figure S8: Time series for the angle of attack of the the (non-covalently bound) photoswitchable inhibitor and the sulfur atom of Cys154 ($\alpha_{S,C_\beta,C_\alpha}$). The shaded area indicates the reactive condition ($110° \leq \alpha_{S,C_\beta,C_\alpha} \leq 130°$)

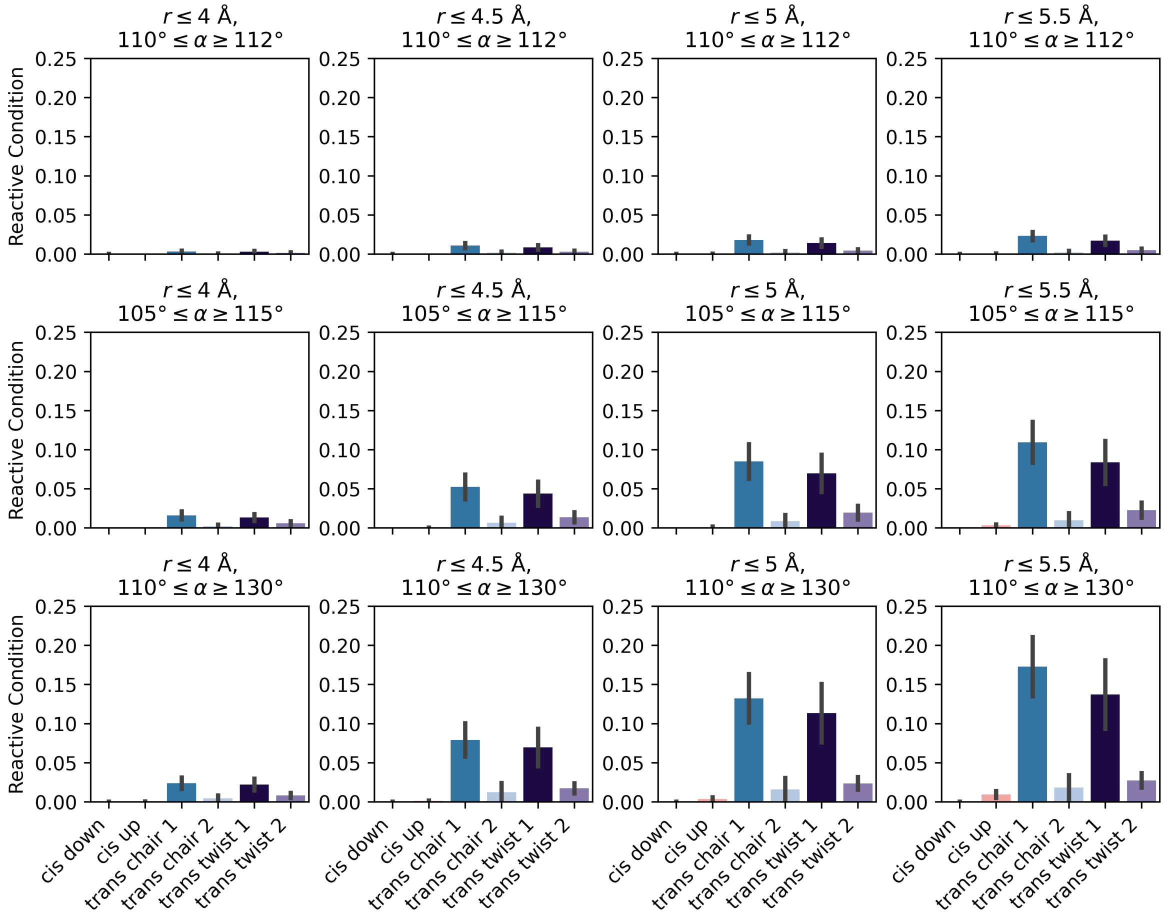


Figure S9: Average number of contacts allowing bond formation between Cys154 and the photoswitchable inhibitor (reactive condition) as shown in Figure 5with different cutoffs for $r_{S,C_\beta}$ and $\alpha_{S,C_\beta,C_\alpha}$. While the absolute values change for different cutoffs, overall trends are consistent.

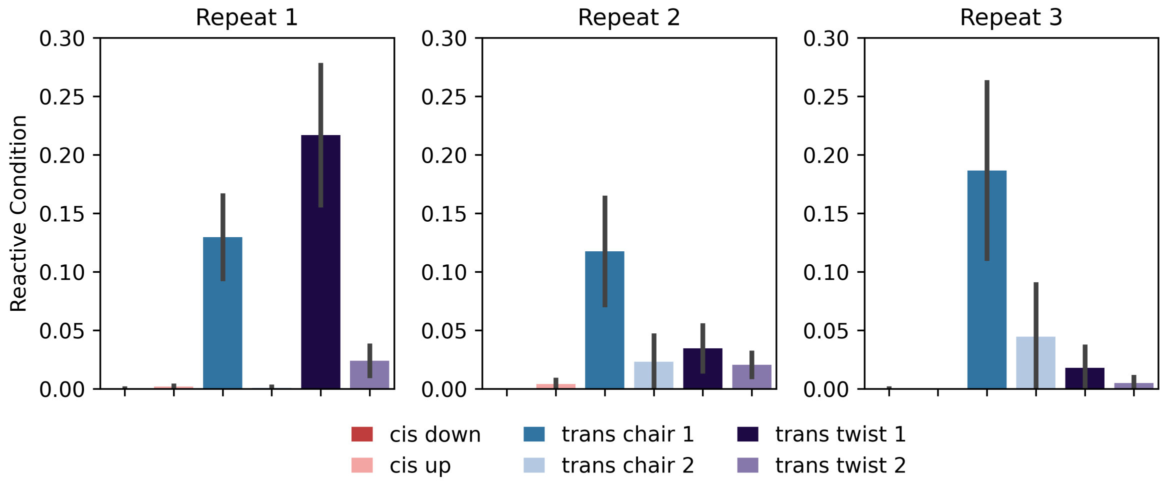


Figure S10: Variability of the reactive conditions between repeats. Average number of contacts allowing bond formation between Cys154 and the photoswitchable inhibitor as shown in Figure 6 for the three repeats, respecively.

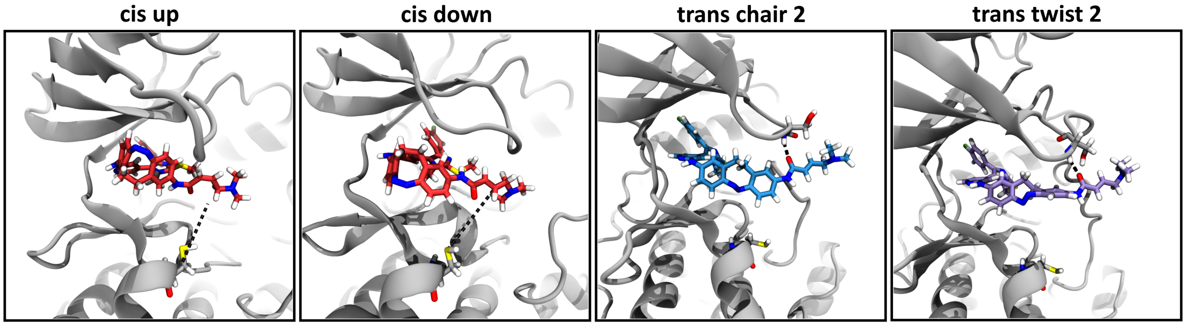


Figure S11: Representative snapshots of the photoswitchable inhibitor bound non-covalently to the ATP-binding pocket of JNK3. For the *cis* up and *cis* down conformation, the distance between the sulfur atom of residue Cys154 and the reactive group of the photoswitchable inhibitor is indicated. For *trans* chair 2 and *trans* twist 2, the hydrogen bond between the peptide oxygen and the backbone of Ser72 is shown.

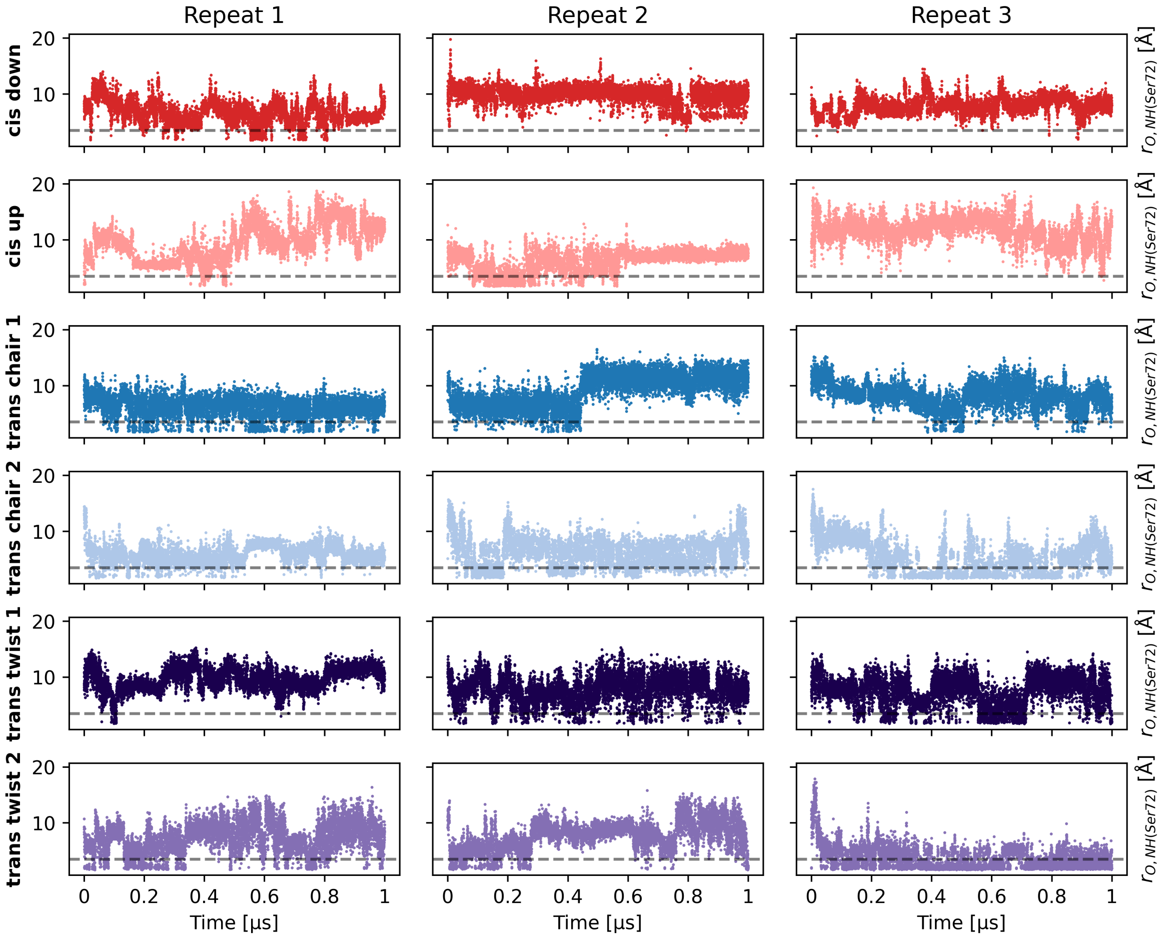


Figure S12: Time series for the distance between the backbone NH of Ser72 and the peptide oxygen of the non-covalently bound photoswitchable inhibitor. The dashed line indicates a value of 3.5 Å.

### S5 Behavior of the photoswitchable inhibitor bound covalently to JNK3

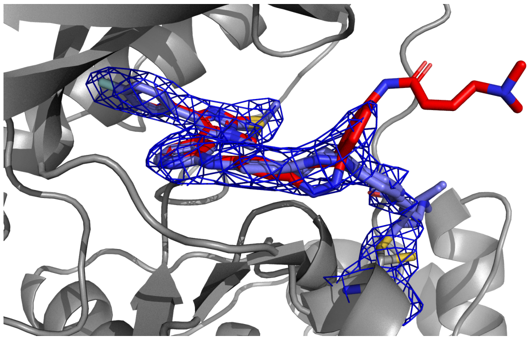

Figure S13: Crystal structure of the photoswitchable inhibitor bound to JNK3. The bent structure is shown in red, while both elongated structures are shown in blue licorice. |2FO|-|FC| refined electron density map contured at $1\sigma$ is shown in blue.

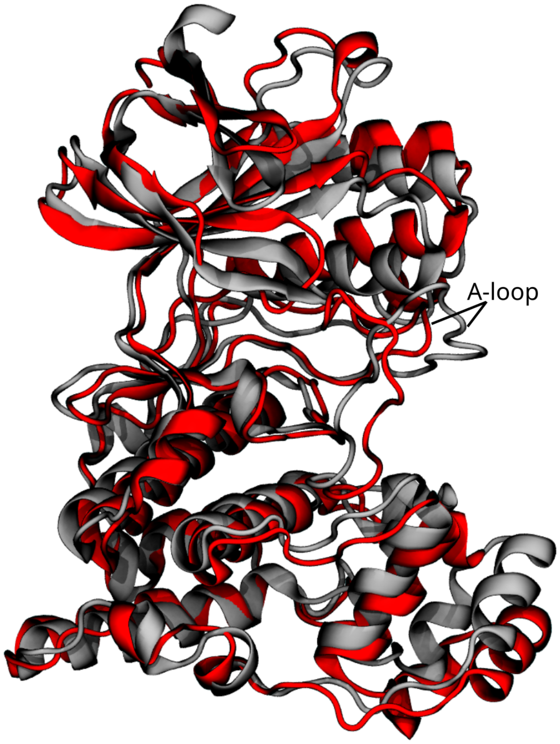


Figure S14: Starting structure of JNK3 for the simulation of the photoswitchable inhibitor in the *cis* up conformation covalently bound shown in red. The crystal structure of JNK3 is shown in grey. While the A-loop is in an intermediate state in the crystal structure, it is in the open state for the simulation with the covalently bound photoswitchable inhibitor in the *cis* up conformation.

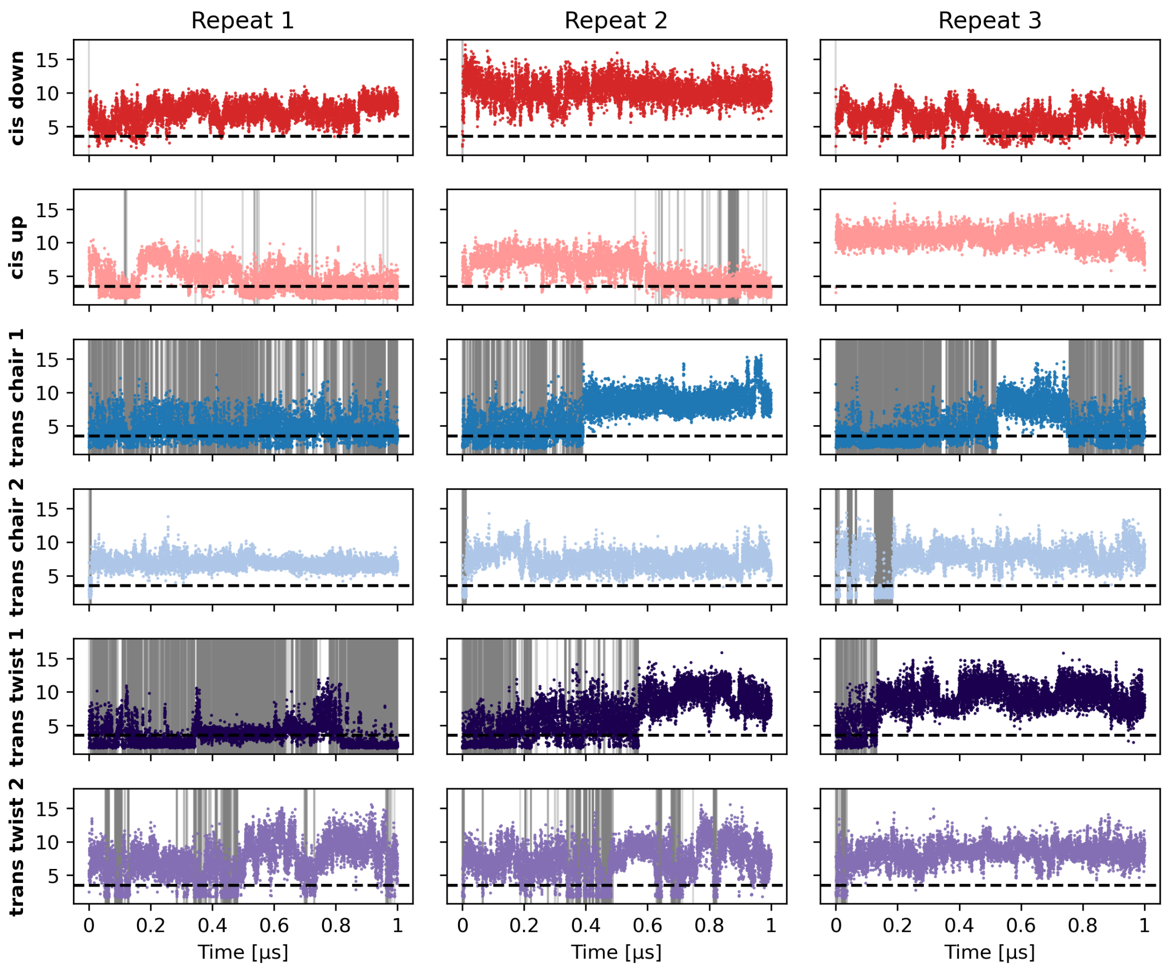


Figure S15: Time series for the distance between the side chain amine of ASN152 and the peptide carbonly oxygen of the non-covalently bound photoswitchable inhibitor. Shown is the minimum distance between the hydrogens of the aminde group and the oxygen atom. The shaded areas indicate frames where the reactive condition is met, while the dashed line indicates a value of 3.5 Å.

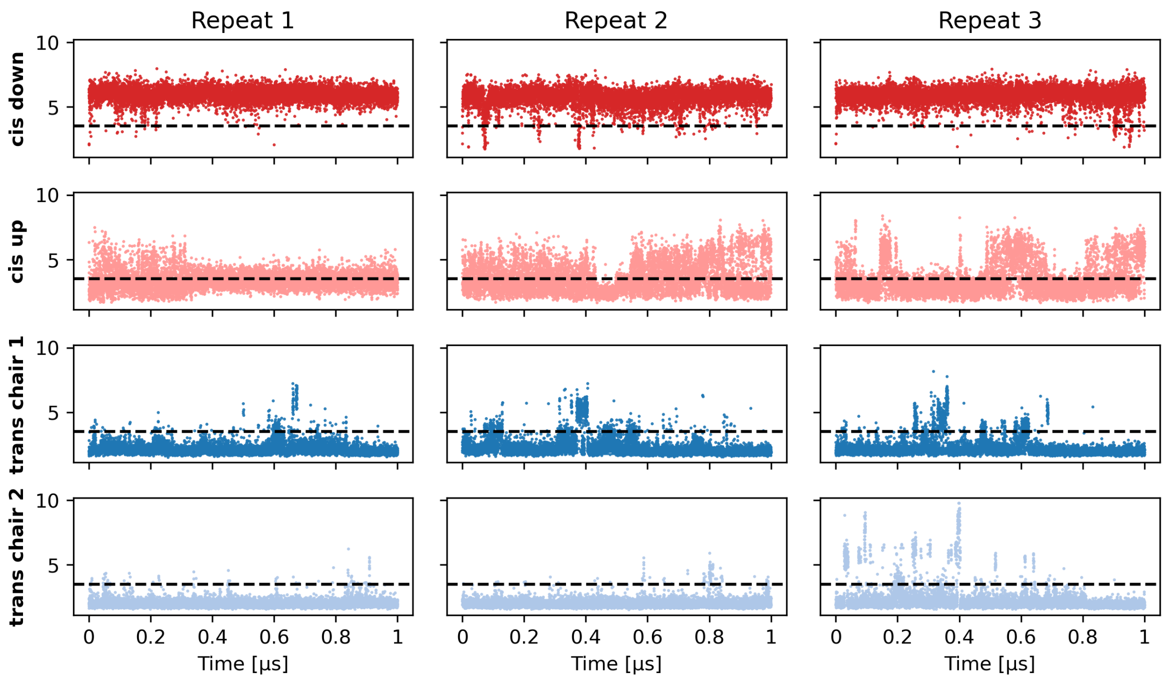


Figure S16: Time series for the distance between the side chain amine of ASN152 and the peptide carbonly oxygen of the covalently bound photoswitchable inhibitor. Shown is the minimum distance between the hydrogens of the aminde group and the oxygen atom. The dashed line indicates a value of 3.5 Å.

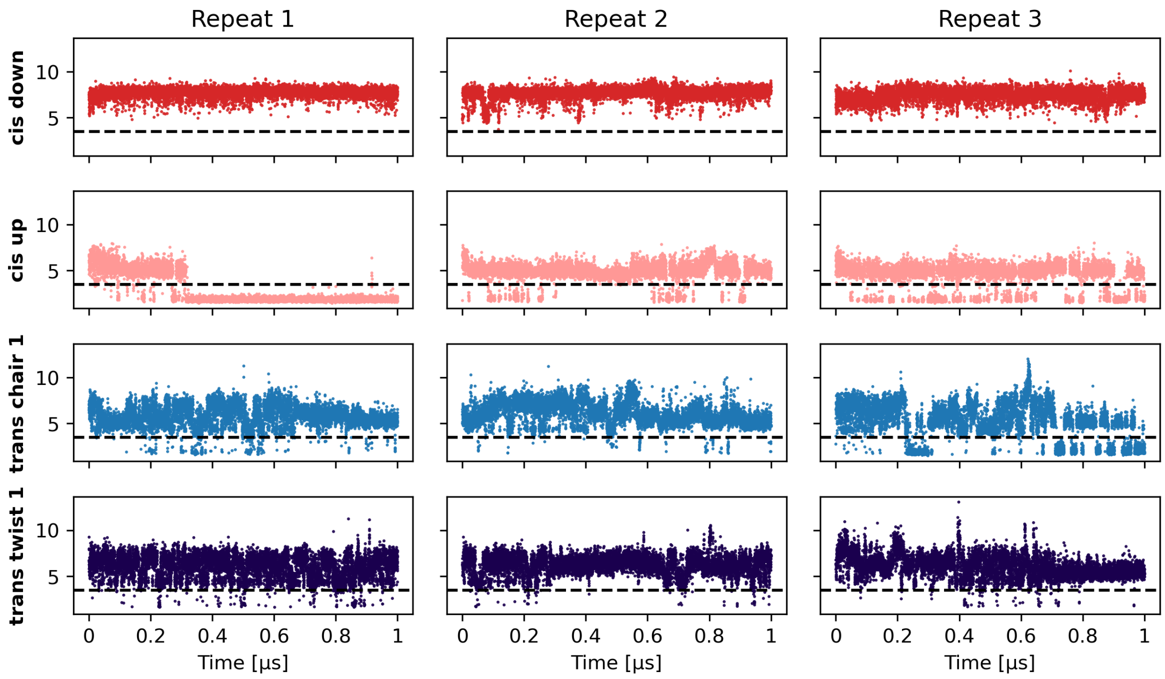


Figure S17: Time series for the distance between the side chain hydroxy group of SER152 and the peptide carbonly oxygen of the covalently bound photoswitchable inhibitor.The dashed line indicates a value of 3.5 Å.

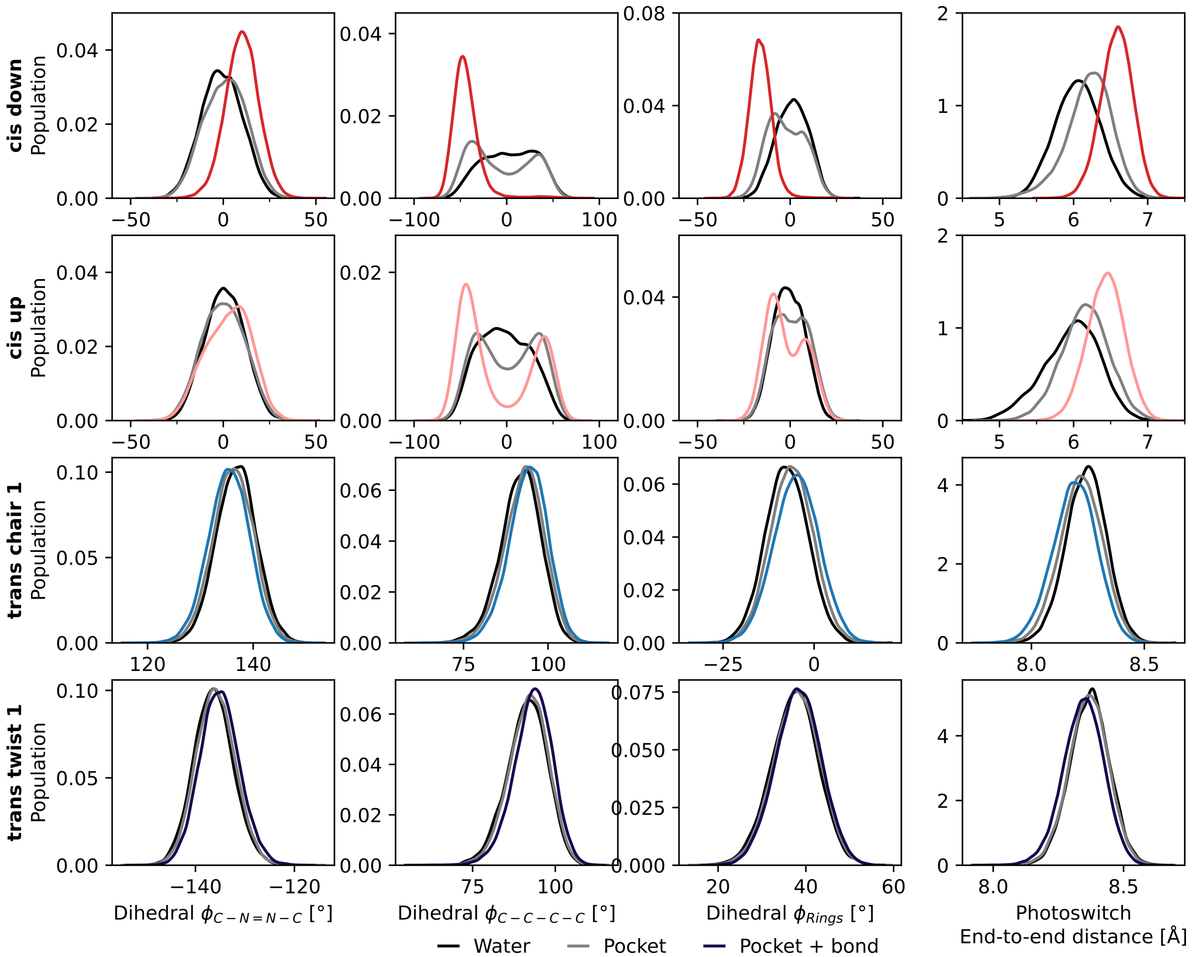


Figure S18: Distributions of the characteristic photoswitch dihedral angles and end-to-end distance for three systems – the photoswitchable inhibitor in water, bound non-covalently (pocket), and covalently (pocket + bond) to JNK3. The dihedral angles are indicated in Figure 3 and the distance in Figure S1.

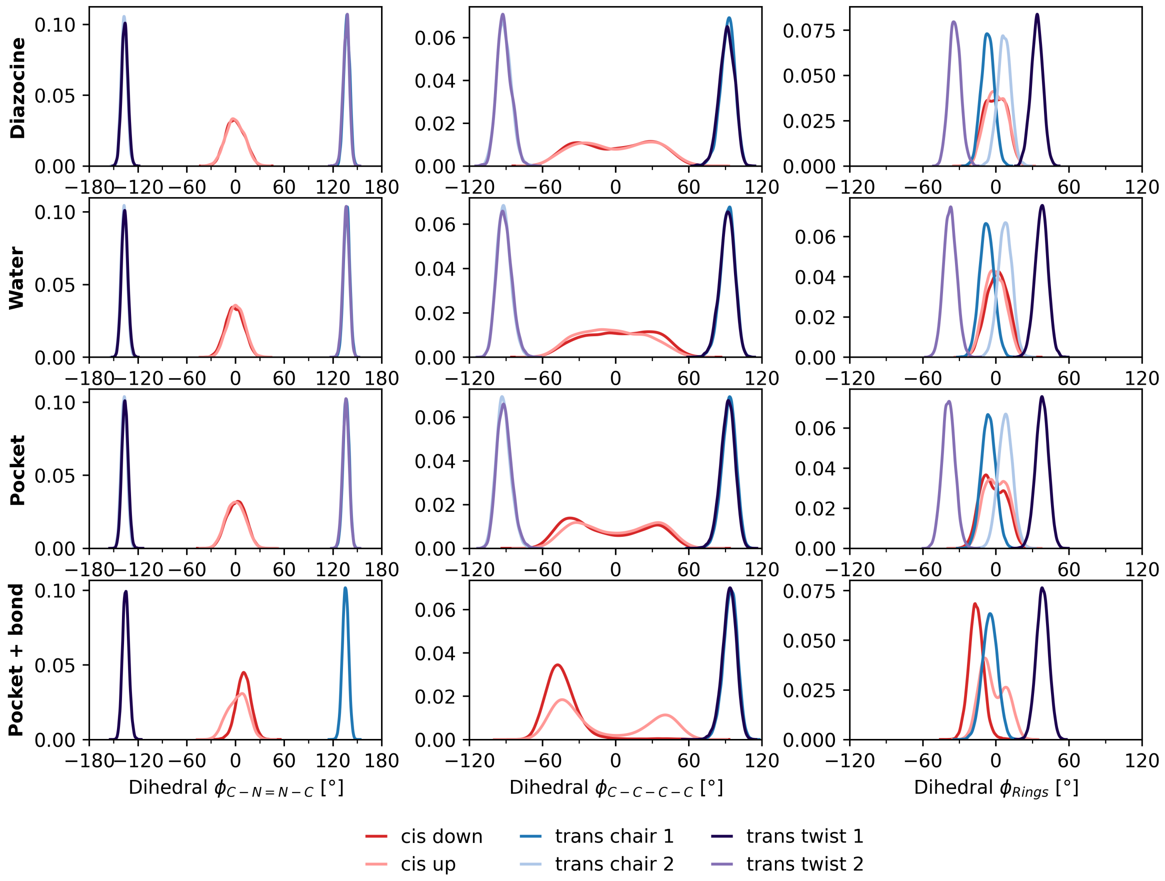


Figure S19: Distributions of the characteristic dihedral angles of the diazocine photoswitch for different conditions (in diazocine, in the photoswitchable inhibitor in water, and bound (non-)covalently to JNK3). The dihedral angles are indicated in Figure 3C.

### S6 Impact of the photoswitchable inhibitor on JNK3 conformation

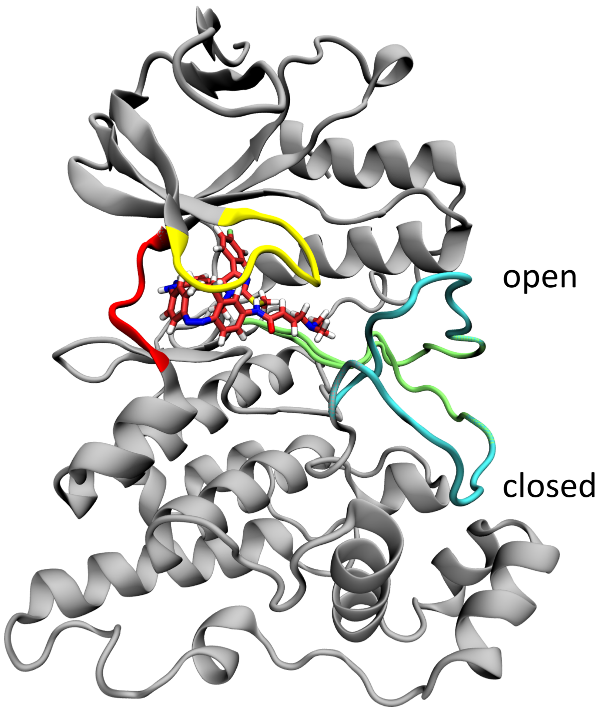


Figure S20: Exemplary snapshots of the A-loop (cyan) in the open and closed conformation. The photoswitchable inhibitor in the *cis* down conformation bound non-covalently to the ATP-binding pocket is shown in red licorice. The following regions are shown in the respective color: G-loop (residues 71-78, yellow), hinge region (residues 147-152, red), A-loop (residues 217-226, cyan), and the activation segment, containing the A-loop (residues 207-226, lime).

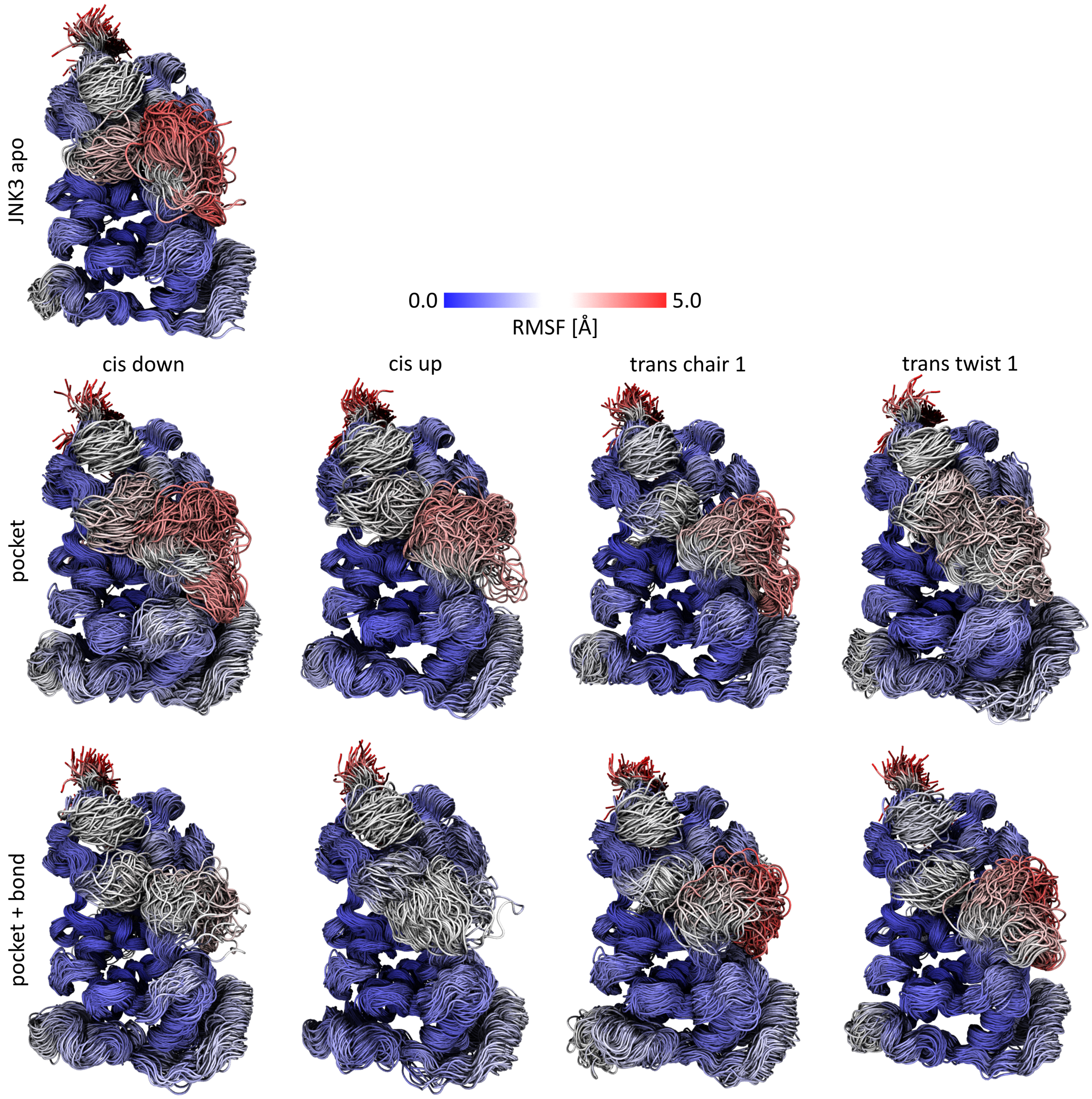


Figure S21: Protein conformations of JNK3 in the apo state and with the photoswitchable inhibitor bound non-covalently (pocket) and covalently (pocket + bond) to the ATP-binding pocket. 300 frames across 3 repeats are shown; the $C_\alpha$ RMSF of JNK3 is indicated by the color.

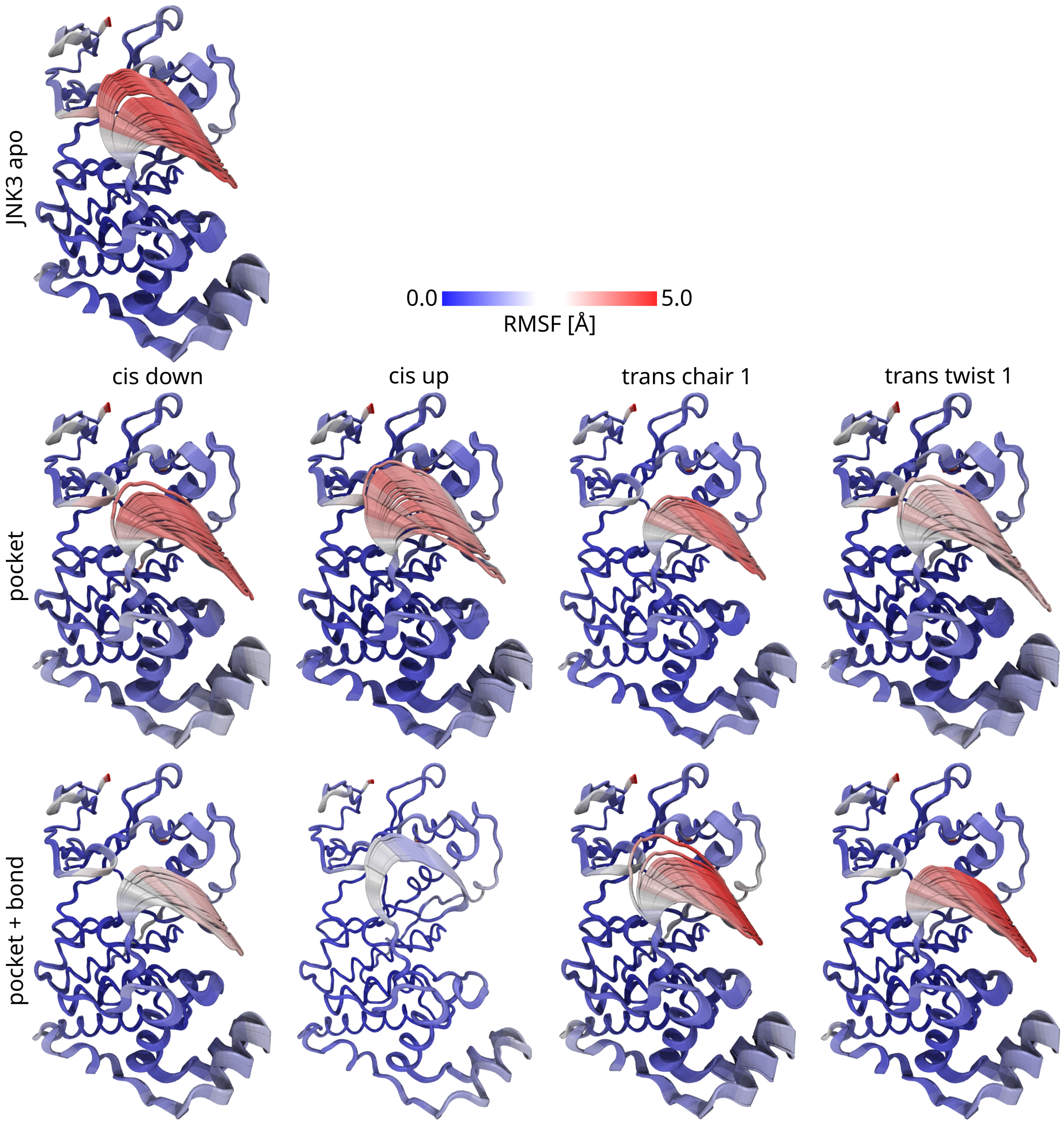


Figure S22: First principle component projected on the protein conformations of JNK3 in the apo state and and with the photoswitchable inhibitor bound non-covalently (pocket) and covalently (pocket + bond) to the ATP-binding pocket. 300 frames across 3 repeats are shown; the $C_\alpha$ RMSF JNK3 is indicated by the color.

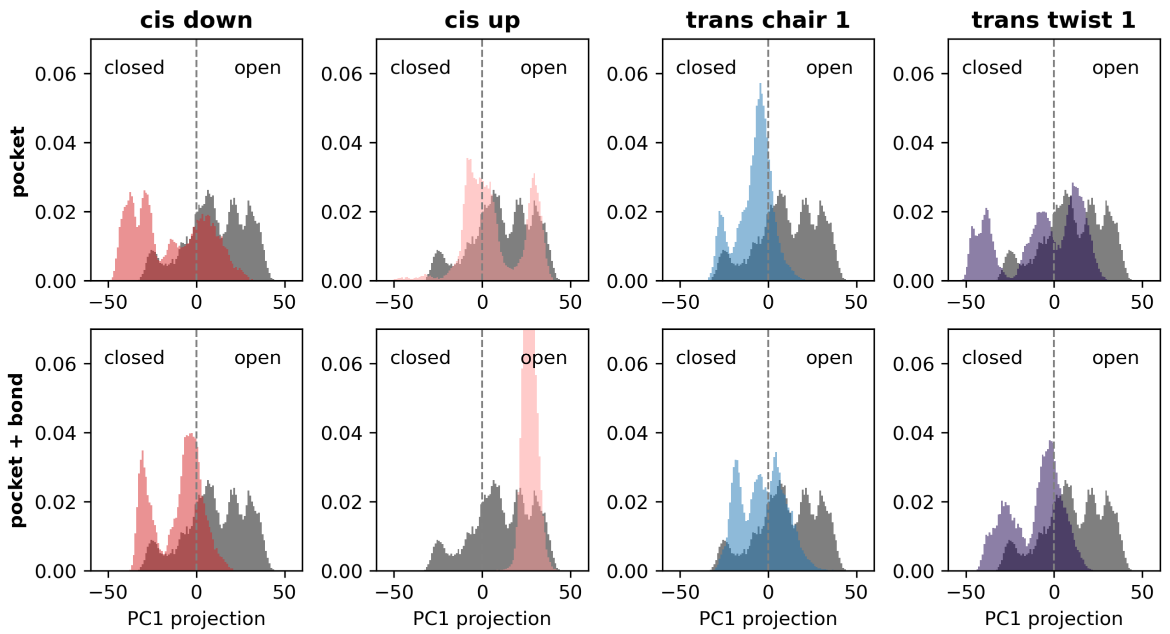


Figure S23: Histograms of the first principle component, which describes the movement of the A-loop for simulations of JNK3 in the apo state and and with the photoswitchable inhibitor bound non-covalently (pocket) and covalently (pocket + bond) to the ATP-binding pocket.

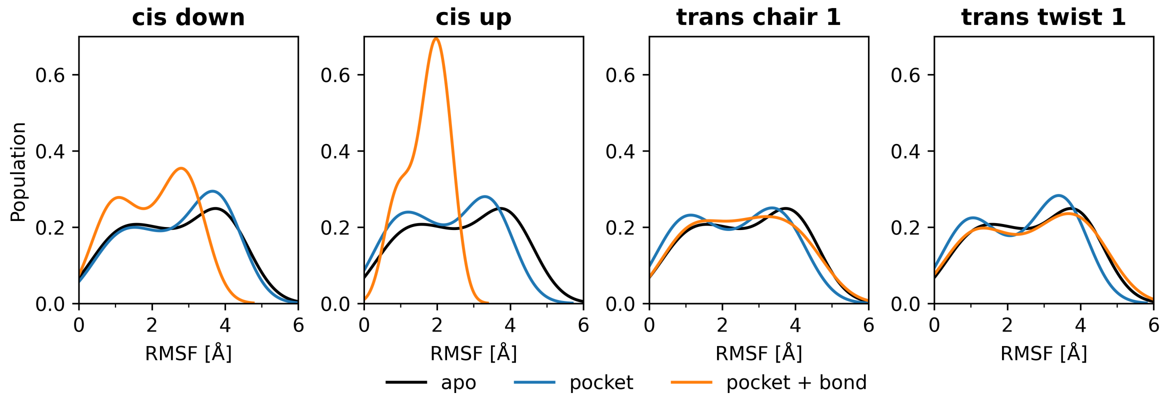


Figure S24: Distribution of the backbone ($C_\alpha$) RMSF of the activation segment of JNK3 (residues 207-226) in the apo state and with the photoswitchable inhibitor bound non-covalently and covalently to the ATP-binding pocket of JNK3.

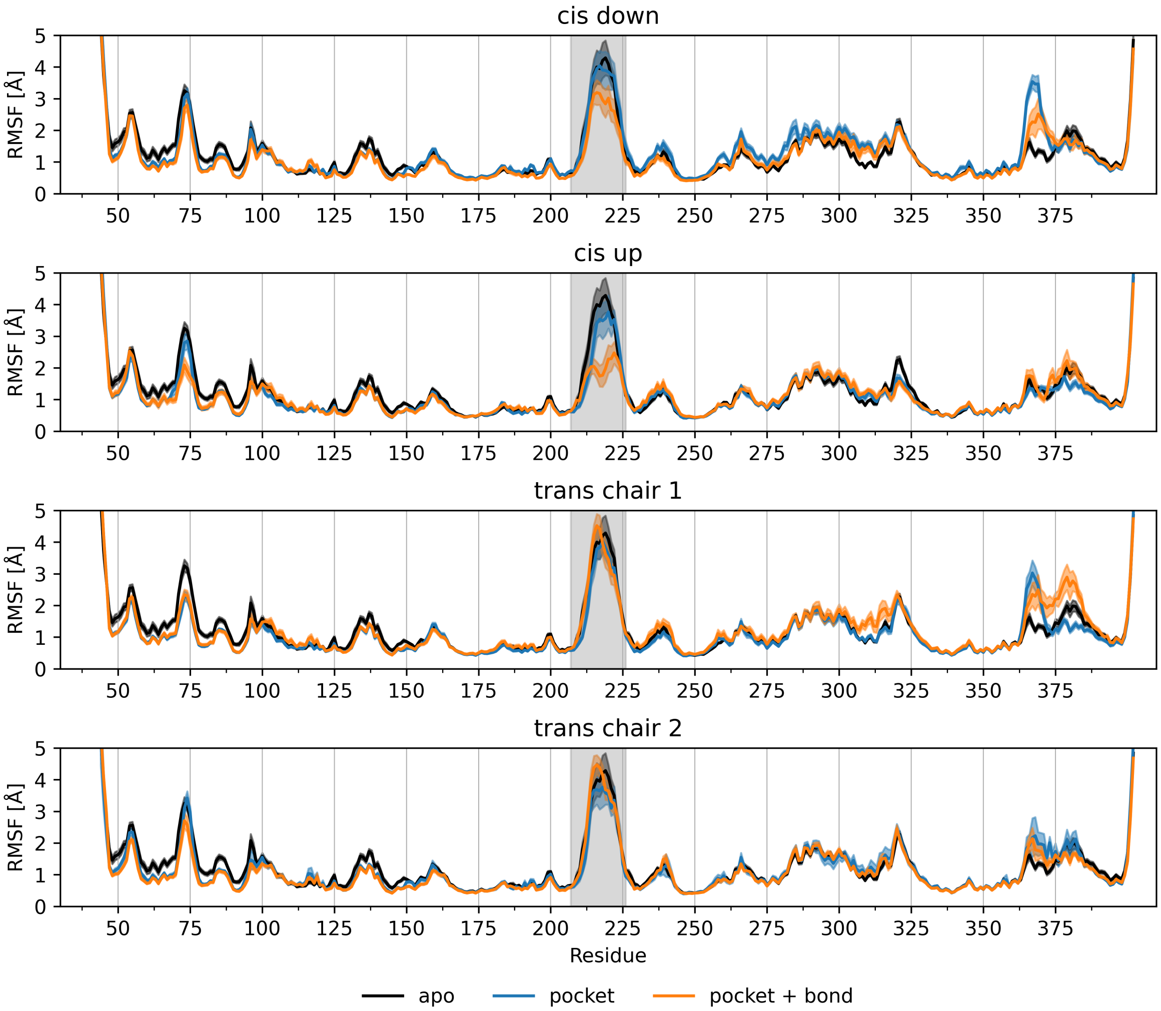


Figure S25: Backbone ($C_\alpha$) RMSF of JNK3 in the apo state and with the photoswitchable inhibitor bound non-covalently and covalently to the ATP-binding pocket of JNK3. The activation segment is shaded in grey.
